# Germline determinants of the prostate tumor genome

**DOI:** 10.1101/2022.11.16.516773

**Authors:** Kathleen E. Houlahan, Jiapei Yuan, Tommer Schwarz, Julie Livingstone, Natalie S. Fox, Weerachai Jaratlerdsiri, Job van Riet, Kodi Taraszka, Natalie Kurganovs, Helen Zhu, Jocelyn Sietsma Penington, Chol-Hee Jung, Takafumi N Yamaguchi, Jue Jiang, Lawrence E Heisler, Richard Jovelin, Susmita G Ramanand, Connor Bell, Edward O’Connor, Shingai B.A. Mutambirwa, Ji-Heui Seo, Anthony J. Costello, Mark M. Pomerantz, Bernard J. Pope, Noah Zaitlen, Amar U. Kishan, Niall M. Corcoran, Robert G. Bristow, Sebastian M. Waszak, Riana M.S. Bornman, Alexander Gusev, Martijn P. Lolkema, Joachim Weischenfeldt, Rayjean J. Hung, Housheng H. He, Vanessa M. Hayes, Bogdan Pasaniuc, Matthew L. Freedman, Christopher M. Hovens, Ram S. Mani, Paul C. Boutros

**Author notes:** Corresponding author at: UCLA Department of Human Genetics, BOX 957088, 57200A South Tower CHS; Los Angeles, CA 90095. Stanford Cancer Institute, Stanford University School of Medicine, CA.

## Abstract

A person’s germline genome strongly influences their risk of developing cancer. Yet the molecular mechanisms linking the host genome to the specific somatic molecular phenotypes of individual cancers are largely unknown. We quantified the relationships between germline polymorphisms and somatic mutational features in prostate cancer. Across 1,991 prostate tumors, we identified 23 co-occurring germline and somatic events in close 2D or 3D spatial genomic proximity, affecting 10 cancer driver genes. These driver quantitative trait loci (dQTLs) overlap active regulatory regions, and shape the tumor epigenome, transcriptome and proteome. Some dQTLs are active in multiple cancer types, and information content analyses imply hundreds of undiscovered dQTLs. Specific dQTLs explain at least 16.7% ancestry-biases in rates of *TMPRSS2-ERG* gene fusions and 67.3% of ancestry-biases in rates of *FOXA1* point mutations. These data reveal extensive influences of common germline variation on somatic mutational landscapes.

## Introduction

Cancers result from the accumulation of genomic and epigenomic aberrations that deregulate normal cellular processes^1,2^. These aberrations can arise from environmental influences, genetic susceptibility or stochastic errors^3^. The exact contribution of each of these three factors to the mutational landscape of any specific tumor is largely unknown, as are the ways in which these factors interact. Varying contributions of these factors and interactions between them result in each individual tumor having a unique mutational composition. This inter-tumoral heterogeneity is a key driver of clinical urgency for precision care.

Of these three factors, the influences of germline genetics on cancer are well-known. About a third of the risk of cancer diagnosis is heritable^4^. Genome-wide association studies (GWAS) have identified hundreds of specific sequence variations associated with risk of diagnosis – predominantly single nucleotide polymorphisms (SNPs)^5–7^. The mechanisms by which germline predisposition loci modulate risk are mostly unknown, but one hypothesis is that they influence somatic mutational evolution. To test this, we focused on prostate cancer: the second most common malignancy in men worldwide^8^, and one of the most heritable. It is estimated that 57% of the variability in prostate cancer diagnosis is explained by genetic factors^4^. Polygenic risk scores (PRS) based on common germline variants can predict risk of a prostate cancer diagnosis for individual men^9,10^. Rare germline variants in DNA damage repair genes or transcription factors like *HOXB13* are associated with both increased risk of diagnosis and increased disease aggression^11–13^. Genetic ancestry is also associated with the somatic landscape of prostate cancer: the *TMPRSS2-ERG* (T2E) fusion occurs less frequently in cancers arising in men of African and Asian ancestry than of European ancestry^14–18^. Localized prostate tumors arising in men who carry deleterious germline *BRCA2* mutations have a somatic mutational profile resembling metastatic castrate-resistant disease^19^. Similarly, specific germline SNPs are associated with *PTEN* deletion^20^ and somatic point mutations in the driver gene *SPOP*^21^. The prostate cancer epigenome is strongly influenced by a patient’s germline genome, with thousands of SNPs influencing methylation status^22,23^, many associated with patient survival and tumor gene expression^23^. Thus, accumulating evidence from rare and common variants and studies of ancestry hint at broad germline-somatic interactions.

We therefore quantified the relationships between germline SNPs and somatic mutational profiles in prostate cancer. We termed SNPs that co-occur with specific prostate cancer driver genes, driver quantitative trait loci (dQTLs). Integrating linear and three-dimensional analysis of DNA structure, we identify 35 dQTLs affecting 10 driver genes in primary localized prostate cancer. Of these, 11 remained statistically significant in a 1,991-patient meta-analysis spanning early onset, primary and metastatic disease. These dQTLs associate with almost every aspect of prostate cancer: methylation, chromatin structure, mRNA abundance, protein abundance and grade at diagnosis. Several affect multiple cancer types. Specific dQTLs associated with somatic *TMPRSS2*-*ERG* fusion and *FOXA1* point mutations explain significant fractions of observed differences in mutation frequencies across ancestry groups. Finally, information content analyses suggest hundreds of undiscovered dQTLs remain, quantifying how the germline genome shapes tumor evolution.

## Results

### Experimental and cohort design

We assembled a discovery cohort of 427 patients with localized prostate cancer, each with whole-genome sequencing (WGS) of blood (mean 39x coverage) and tumor (mean 64x coverage)^24–27^. All patients had localized disease at diagnosis and were treated by image-guided radiotherapy or surgery with curative intent. All discovery cohort samples were treatment-naïve and macro-dissected by a genitourinary pathologist to obtain 60%+ tumor cellularity, as verified computationally (**Supplementary Table 1**)^28^. Median follow-up was 7.7 years: clinical and molecular data, including indications of germline variants in homologous repair genes, mismatch repair genes and *HOXB13*, are in **Supplementary Table 1**. Patients were of European ancestry and identity-by-state clustering did not reveal population substructure (**Supplementary Figure 1a**). Sequencing data were uniformly processed from read level using benchmarked pipelines^29,30^. We identified 17 somatic drivers occurring in at least 5% of patients based on enrichment over the local background mutational rate and with literature support and focused our analyses on these (range: 5.1-57.3%; **Supplementary Figure 1b**). These comprised 14 copy number aberrations (CNAs), two single nucleotide variants (SNVs) and the fusion of *TMPRSS2* and *ERG* (T2E) ^28^. CNAs were annotated as present in all tumor cells (*i*.*e*., clonal; referred to as trunk) *vs*. a subset of tumor cells (*i*.*e*., subclonal; referred to as branch).

We sought to determine whether individual germline SNPs were associated with specific driver mutations; we termed these driver quantitative trait loci (dQTLs). A fully powered genome-wide discovery would require many thousands of patients with tumor whole-genome sequencing. To enrich for dQTLs, we therefore created three complementary, biologically motivated approaches (**Figure 1a**). First, we tested if germline SNPs associated with risk of diagnosis in prostate-cancer GWAS studies were dQTLs. Second, we identified local dQTLs: regions in close proximity to each somatic driver based on linear DNA sequence. Third, we exploited knowledge of three-dimensional DNA structure to identify spatial local dQTLs. Altogether we evaluated 5,516 independent SNPs against one of 17 somatic drivers.

**Figure 1.**
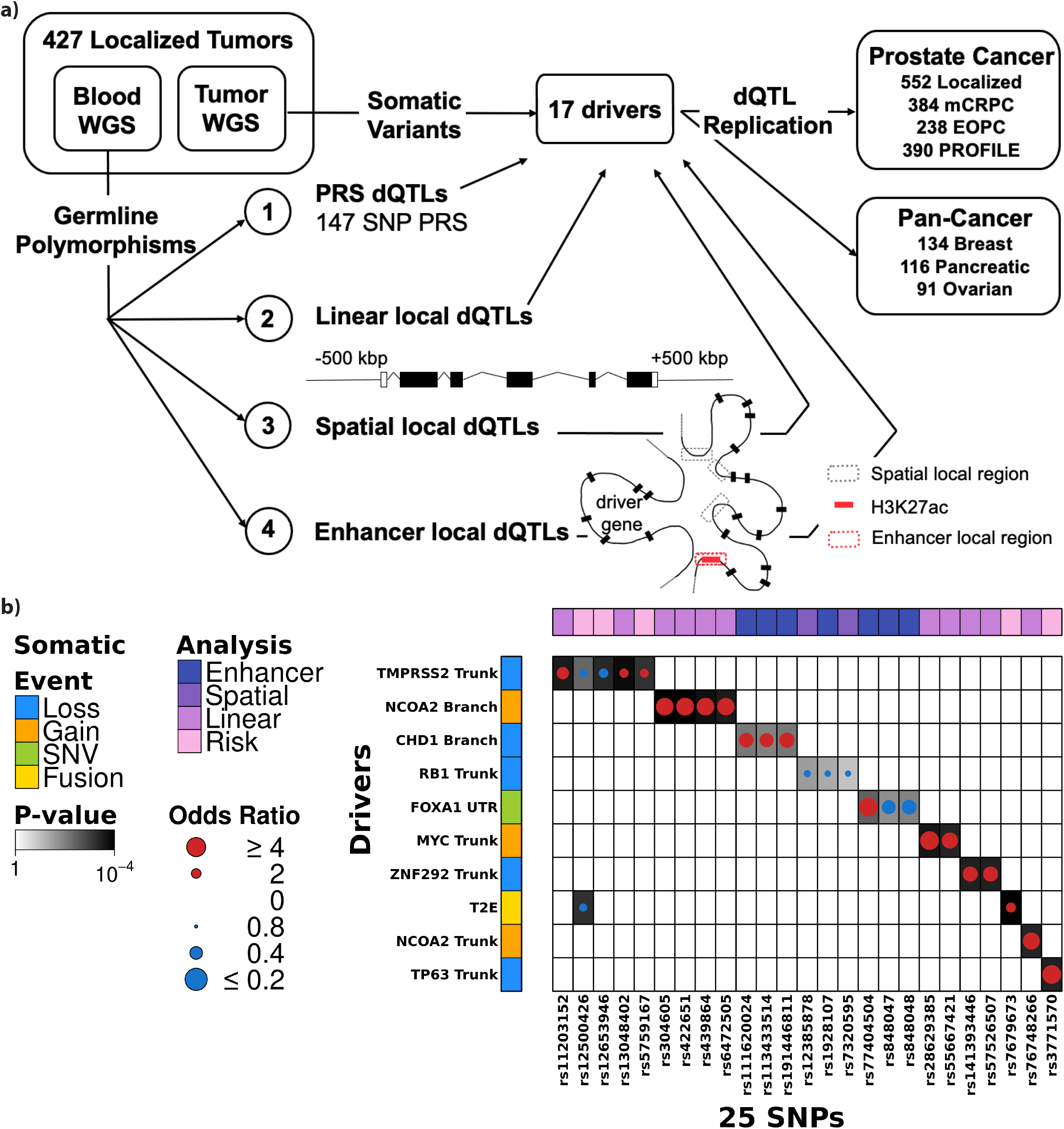
dQTLs bias somatic mutational landscape. **a)** Schematic of dQTL detection. First, 147 SNPs from the polygenic risk score proposed by Schumacher *et al*. ^9^ were interrogated for their association with 17 somatic drivers. Second, we identified linear local dQTLs by interrogating SNPs ±500kbp around the driver gene. Third, we identified spatial local dQTLs by interrogating SNPs that interacted with each driver gene in 3D space, outside of the linear gene region. Spatial local regions were defined using RNA Pol-II ChIA-PET profiling in LNCaP, DU145, VCaP and RWPE-1 cell lines and RAD21 ChIA-PET in LNCaP and DU145 cell lines. Finally, we identified enhancer local dQTLs by interrogating SNPs in enhancer regions that interacted with the driver gene. Enhancer regions were defined using H3K27ac HiChIP profiling in LNCaP cell lines. All discovered dQTLs were tested for replication in six replication cohorts. **b)** Summary of 26 discovery dQTLs involving 25 unique variants. Dot size and color indicates magnitude and direction of ORs between SNP, x-axis and somatic driver, y-axis. Background shading indicates p-values. Covariate on left indicates the type of somatic mutation while covariate along the top indicates the analysis in which the dQTL was discovered.

For replication we compiled a 552-patient cohort of tumors arising in men of European descent from The Cancer Genome Atlas (TCGA) ^31^ supplemented by 140 primary prostate cancers with blood and tumor whole genome sequencing analyzed identically to the discovery cohort (**Supplementary Table 1**)^28^. Finally, to assess dQTL generalizability, we analyzed cancer types in the Pan-Cancer Analysis of Whole Genomes (PCAWG) cohort with at least 90 individuals of European descent (*i*.*e*., breast, ovarian and pancreatic)^32^. **Supplementary Table 1** summarizes all cohorts evaluated.

As a positive control, we first replicated previously reported SNP associations. Two SNPs associated with T2E were replicated: rs16901979 (OR = 0.50; P = 3.90 × 10^−2^; **Supplementary Figure 1c**) and rs1859962 (OR = 1.52; P = 5.05 × 10^−3^; **Supplementary Figure 1d**)^33^. Two SNPs in *HSD3B1* associated with overall survival in advanced prostate cancer^34^ showed trend associations with clinical relapse (P_rs1856888_ = 0.11; P_rs1047303_ = 0.18; **Supplementary Figure 1e-f)** and with tumor extent at diagnosis (P_rs1856888_ = 0.029; P_rs1047303_ = 0.091; **Supplementary Figure 1g-h)**. SNPs reported to be associated with *PTEN* loss^20^ and *SPOP* point mutations were not replicated in this cohort^21^. Finally, the observation in melanoma that SNPs in *APOE* were associated with metastasis-free survival was generalized to prostate cancer (P = 0.027; **Supplementary Figure 1i**)^35^. Tumors with the APOE2 genotype had a significantly higher burden of genomic rearrangements (GRs) than APOE4 tumors (OR = 0.45; P = 0.05; **Supplementary Figure 1j**). These positive controls confirm our patient cohorts replicate known germline-somatic associations but highlight the potential for false negatives at this statistical power as well as false positives in published candidate gene approaches.

### Risk variants associated with somatic drivers in prostate cancer

To identify germline SNPs associated with somatic drivers, we first considered risk alleles used in a polygenic risk score (PRS) derived from 147 variants^9^ (**Figure 1a**). Of the 134 individual risk SNPs with a minor allele frequency (MAF) > 0.05 in the discovery cohort, six were associated with one or more somatic driver mutations (logistic regression; Benjamini-Hochberg (BH) FDR < 0.1; labelled in light pink in **Figure 1b; Supplementary Figure 2a; Supplementary Table 2&3**). rs12500426, was associated with both loss of *TMPRSS2* and T2E gene fusion, as expected (OR = 0.60 & 0.59; BH FDR = 0.095 & 0.027, respectively; **Figure 1b**). To control for index event bias, we confirmed the six dQTLs after correcting for ISUP grade group, T category and PSA levels (P < 7.8×10^−3^; **Supplementary Table 2**). We replicated previous reports of rs7679673 (OR = 1.94; BH FDR = 0.011) and rs12653946 (OR = 0.53; BH FDR = 0.032) association with ERG status^36^ (**Figure 1b**). To increase statistical power, we grouped somatic driver by pathways: *ETS* fusions (*i*.*e*., fusions in any *ETS* gene), cell cycle (loss of *CDKN1B* or *RB1*) and AR signaling (loss of *NKX3-1*, SNVs in *FOXA1* or gain of *NCOA2*). No additional pathway-based dQTLs were identified.

Finally, we interrogated if the *HOXB13* G84E variant was associated with risk of acquiring any of the 17 somatic drivers. Because *HOXB13* G84E is not common (MAF = 0.0024), we combined the 427 patients in the discovery cohort with the 552 patients in the replication cohort. In the 15 patients with *HOXB13* G84E (**Supplementary Table 1**), there was a nominal association with T2E (OR = 0.27; P = 0.046) which did not survive multiple hypothesis testing (**Supplementary Table 2**). Cohorts with more *HOXB13* G84E carriers will be required to robustly assess if *HOXB13* G84E is a dQTL.

### Local dQTLs bias somatic drivers in prostate cancer

The association of individual risk alleles with somatic mutations suggested that specific dQTLs might influence the mutational and evolutionary diversity of localized prostate cancer^37^. We evaluated common SNPs (MAF > 0.05) in “close proximity” to somatic driver mutations using three different definitions of “proximity” (**Figure 1a**). First, we defined proximity based on the primary DNA sequence and considered germline variants within ±500 kbp of the somatic event boundaries. This distance threshold was selected through sensitivity analysis (**Supplementary Figure 2b**). The 17 somatic drivers were each compared to 1,332-11,618 germline SNPs (median = 2,279, haplotype blocks = 80-1,379; median haplotype block size = 7 SNPs; **Supplementary Figure 2c**). After controlling for population structure and somatic mutation burden, 20 local dQTLs were identified in 11 haplotype blocks, involving five drivers (logistic regression; Bonferroni α=0.1 per driver; P_unadjusted_ < 3.7×10^−4^; OR > 1.8; **Figure 1b**; **Supplementary Table 3&4**). We selected a tag dQTL – *i*.*e*., one SNP to represent each haplotype block – based on minimum p-value. A subset of patients in our discovery cohort (n=325/427) had additional CNA profiling using orthogonal array-based platforms, and all 11 CNA tag SNPs were verified by this independent technology (**Supplementary Figure 2d**).

Second, we defined proximity to the somatic event based on DNA secondary structure (**Figure 1a**). Spatial local dQTLs were defined based on RNA polymerase II ChIA-PET in LNCaP, DU145 and VCaP prostate cancer cells and RWPE-1 benign prostate epithelial cells ^38^, along with RAD21 ChIA-PET in LNCaP and DU145 cell lines ^39^. We identified regions outside the linear local boundaries that interacted with the event region in at least two of four cell lines. Each of the 17 somatic drivers was evaluated for associations with 7-101 SNPs in this step (median = 32; haplotype blocks = 2-16; median haplotype block size = 3 SNPs; **Supplementary Figure 2e**). Two dQTLs associated with clonal (trunk) loss of *RB1* were discovered (logistic regression; Bonferroni α=0.1 per driver; P_unadjusted_ < 2.35×10^−2^; OR > 1.47; **Figure 1b; Supplementary Table 3&4**), and both verified using array-based CNAs (**Supplementary Figure 2f**).

Finally, to further explore dQTLs in three-dimensional space, we considered proximity as defined by interacting enhancers identified *via* HiChIP H3K27ac profiling in LNCaP cell lines (**Figure 1a**). We identified anchor regions outside of gene boundaries whose associated anchor fell within the driver gene of interest (see **Methods**). The 17 somatic drivers were evaluated for associations with 0-1,059 SNPs (median = 35; haplotype blocks = 0-81; median haplotype block size = 5 SNPs; **Supplementary Figure 2g**). We identified 11 dQTLs involving seven haplotype blocks and three somatic drivers (logistic regression; Bonferroni α=0.1 per driver; P_unadjusted_ < 1.27×10^−2^; OR > 1.50; **Figure 1b; Supplementary Table 3&4**). We verified 3/4 candidate CNA dQTLs using array-based data (3 dQTLs associated with SNVs in *FOXA1* 3’ UTR were not measured on the array platform used; **Supplementary Figure 2h**).

### dQTLs affect multiple drivers and cancer types

We thus identified 26 tag dQTLs involving 25 unique loci using four strategies: risk dQTLs, linear local dQTLs, spatial local dQTLs and enhancer local dQTLs (**Figure 1a**). Despite being significantly under-powered, 16/26 showed consistent effect-sizes in our replication cohort (*i*.*e*., sign(log(OR_discovery_)) = sign(log(OR_replication_)); **Figure 2a**) and four statistically replicated (BH FDR < 0.1). These four were rs11203152 with loss of *TMPRSS2* (a proxy for T2E status), rs141393446 with loss of *ZNF292* and both rs848047 and rs848048 with SNVs in 3’ UTR of *FOXA1* (**Supplementary Figure 3a-h**). Next, we investigated dQTL replication in other cancer types, focusing on ovarian, breast and pancreatic cancers from PCAWG^32^. We tested only somatic drivers with mutation frequencies ≥ 5% in each cancer type (*i*.*e*., 20/26 tag dQTLs). Of these 20, 14 showed consistent effect sizes in breast, ovarian or pancreatic cancers (**Supplementary Figure 3i-k)**. The association between rs76748266 and gain of *NCOA2* replicated in pancreatic cancer (OR_pancreatic_ = 6.47; BH FDR_pancreatic_ = 1.56 × 10^−2^; **Supplementary Figure 3l-m)** and the association between rs11203152 with loss of *TMPRSS2* was nominally significant in ovarian cancer but did not survive multiple hypothesis testing correction (OR_ovarian_ = 4.87; BH FDR_ovarian_ = 0.11; **Supplementary Figure 3n). Supplementary Table 5** includes the summary statistics for dQTLs across discovery and replication cohorts and **Supplementary Table 1** summarizes the cohorts evaluated. Thus, a subset of dQTLs affect multiple cancer types. Prostate cancer genomic studies have identified mutually exclusive and co-occurring somatic mutations^28^. We therefore sought to identify local dQTLs that show associations with distal driver genes. Focusing on the 16 dQTLs with consistent ORs in the replication cohort, we screened each tag SNP against all 17 somatic drivers in a candidate analysis. This identified nine candidate distal dQTLs (BH FDR < 0.1; **Supplementary Figure 3o**; **Supplementary Table 3&6**), seven of which showed concordant ORs in our replication cohort (**Supplementary Figure 3p**). Next, we investigated if dQTLs were associated with chromothripsis, a mechanism that can simultaneously disrupt multiple driver genes^40^, but did not find any associated dQTLs (BH FDR > 0.40; **Supplementary Table 3**). Integrating all our results, we discovered 35 dQTLs involving 25 tag SNPs and 10 somatic drivers (**Figure 2b**). Two thirds showed consistent effect-sizes in our replication cohort (Fold Change (FC) = 1.33, P = 0.028; n = 10,000; permutation test; **Supplementary Figure 3q**), and five replicated (BH FDR < 0.1) in at least one cancer type.

**Figure 2.**
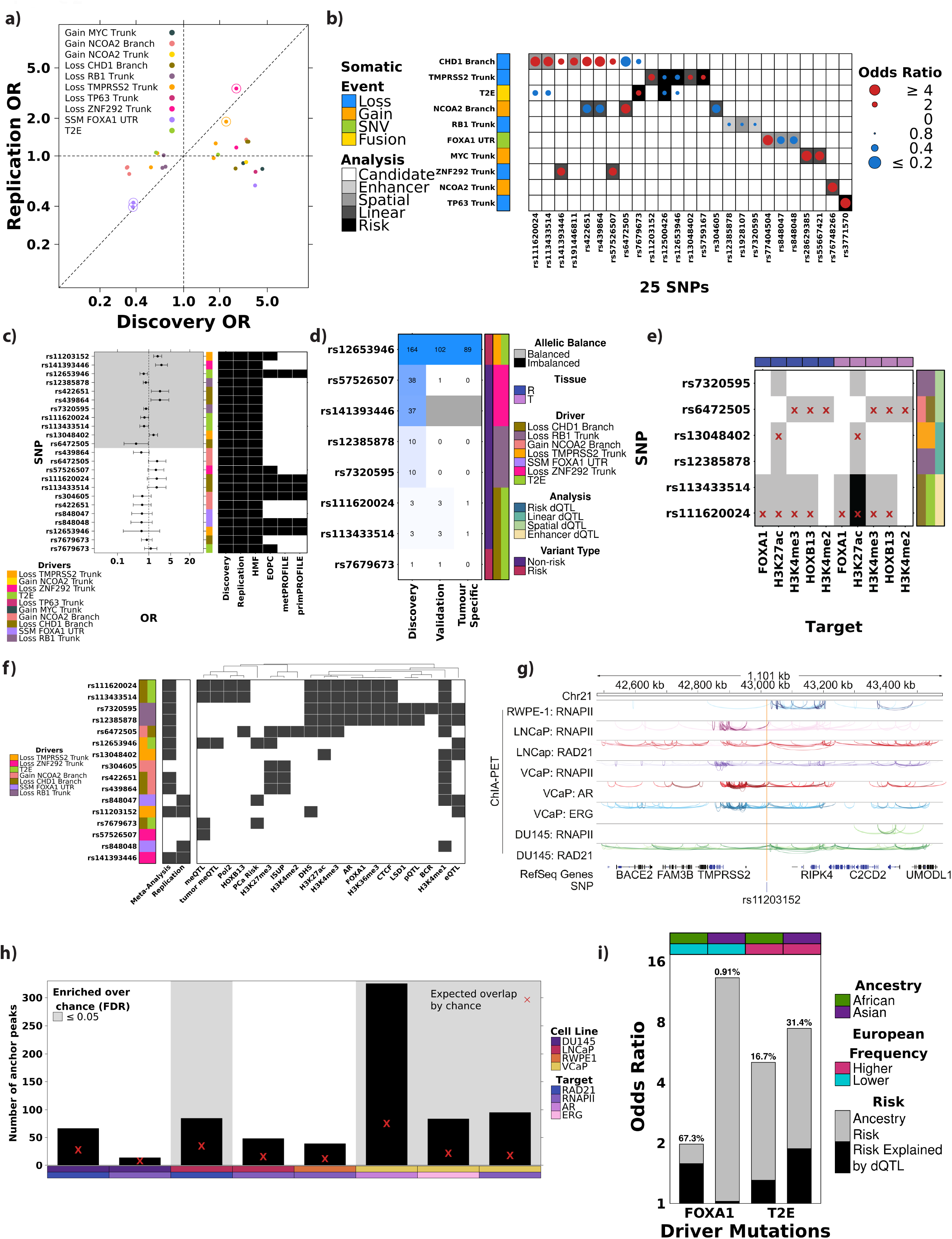
Characterization of dQTLs. **a)** Comparison of ORs in discovery, x-axis, *vs*. replication, y-axis, cohort of tag dQTLs. Horizontal and vertical dotted lines represent OR = 1 and diagonal line represents y=x. Halo around points indicates BH FDR < 0.1 in replication cohort. Dot color indicates the associated somatic driver. **b)** Summary of all 35 dQTLs involving 25 unique variants. Dot size and color indicates magnitude and direction of ORs between SNP, x-axis and somatic driver, y-axis. Background shading indicates strategy dQTL was discovered with. Covariate on left indicates the type of somatic mutation. **c)** Forest plot shows the OR and 95% confidence interval for dQTL associations from a meta-analysis of 2,019 prostate tumors. The grey shading indicates an BH FDR < 0.1. The covariate in the middle indicates the driver mutation the SNP is associated with. The heatmap on the right indicates which cohorts were included in the meta-analysis. **d)** A subset of dQTLs were associated with changes in tumor methylation. Heatmap indicates the number of methylation probes each variant, x-axis, was associated with in the discovery and replication TCGA cohort, y-axis. The third column indicates the number of replicated meQTLs that were tumor specific. The covariate on the right indicates if the variant is a risk variant and what somatic driver it is associated with. **e)** dQTL variants (x-axis) overlap with histone modification and transcription factor binding sites (y-axis). Grey shading indicates overlap with allelic balanced ChIP-Seq peak while black indicates overlap with allelic imbalanced ChIP-Seq peak. Red X indicates overlapping SNP is tag SNP. Covariate along the top indicates the tissue while the covariate along the right indicates if the SNP is a literature reported risk SNP and what somatic driver it was associated with. **f)** Summary of molecular and clinical characterization of dQTLs. Grey indicates dQTL was association with methylation (meQTL), RNA abundance (eQTL), protein abundance (pQTL), transcription factor binding, histone modification, ISUP grade group, biochemical recurrence (BCR) or risk of prostate cancer diagnosis (PCa Risk). Middle heatmap shows if dQTL replicated in meta-analysis or the replication cohort. Covariate on the left illustrates the somatic driver the dQTL is associated with. **g)** rs11203152 located within regulatory dense region. Tracks show chromatin looping anchored by RNA Polymerase II (RNAPII), RAD21, AR or ERG in RWPE-1, LNCaP, VCaP or DU145 cell lines. **h)** The number of chromatin loops was more than expected by chance in LNCaP and VCaP cell lines. Barplots shows number of anchors within 1 Mbp of rs11203152. Covariate along the bottom indicates cell line and target while the background shading indicates of the enrichment was more than expected by chance (BH FDR < 0.05). The red X indicates the expected number of chromatin loop anchors based on 100,000 randomly sampled, equally sized regions. **i)** dQTLs may, in part, explain differences in somatic mutation frequencies across ancestries. Barplot shows the risk (y-axis) of acquiring a *FOXA1* SNV or T2E based African (green) or Asian (purple) ancestry compared to European ancestry (x-axis). The estimated percent of this risk explained by rs848048 (*FOXA1*) or rs11203152 (T2E) is shown in black and indicated above the bar. The covariate along the top indicates if the bar represents African (green) or Asian (purple) descent individuals and if the somatic mutation is observed more (pink) or less (teal) frequently in European-descent men compared to African or Asian-descent.

### dQTLs generalize to other types of prostate cancer

To extend these results to other forms of prostate cancer and increase our replication power, we considered early-onset (EOPC; diagnosis < 55 years) and metastatic prostate tumors (**Supplementary Table 1**). We conducted a meta-analysis across 1,991 European descent prostate tumors, including the discovery and replication cohorts as well as 238 EOPC tumors^41^, 384 metastatic castrate resistant prostate tumors^42^, and 91 metastatic and 299 localized prostate tumors from the PROFILE cohort^43^. We focused on 23 dQTLs that showed concordant ORs in the discovery and replication cohorts (henceforth termed concordant dQTLs; **Supplementary Figure 3q**). Not all dQTLs could be tested in each cohort because of limitations in sequencing protocols – *e*.*g*., PROFILE used targeted sequencing – and limited power due to differences in driver mutation recurrence rates across disease stages. **Figure 2c** indicates in which cohorts each dQTL was tested. We identified 11 statistically replicated dQTLs (BH FDR < 0.1; **Figure 2c**; **Supplementary Table 5**) across these 1,991 patients. Thus, dQTLs can generalize across stages of prostate cancer.

### Local dQTLs modulate the tumor epigenome

Deregulation of tumor methylation is one mechanism by which the germline genome influences cancer risk ^22,23^. We investigated if dQTL tag SNPs were associated with methylation changes in tumor tissue (**Supplementary Figure 4a**). We focused on the 23 concordant tag dQTLs that showed consistent ORs in the replication cohort (**Supplementary Figure 3q**) and conducted a candidate local methylation quantitative trait loci (meQTL) analysis within ±500 kbp of dQTL tag SNPs. We used array-based methylomes from 226 patients in the discovery cohort and 412 patients in the replication cohort, along with 47 profiles of histologically non-malignant reference prostate tissue (**Supplementary Table 1**). We identified 266 local meQTLs involving eight dQTLs (|β_discovery_| > 0.041; BH FDR_discovery_ < 0.1). Our replication cohort had genotyping of 221/266 local meQTLs, and 110 replicated (|β_replication_| > 0.039; BH FDR_replication_ < 0.1; **Figure 2d Supplementary Figure 4b**; **Supplementary Table 7**). Three SNPs, rs12653946 associated with T2E and clonal loss of *TMPRSS2*, and rs111620024 and rs113433514 associated with T2E and subclonal loss of *CHD1*, were involved in tumor-specific meQTLs: they were associated with methylation changes in tumor but not normal cells ^23^ (|β_tumor_| > 0.12; BH FDR_tumor_ < 8.50×10^−2^; |β_reference_| < 0.63; BH FDR_reference_ > 0.12). There were significantly more methylation probes associated with dQTLs than expected by chance (P < 10^−4^; observed meQTLs = 110; expected meQTLs = 10; permutation test), but not more dQTLs exhibiting meQTL behavior (P = 0.42; permutation n = 1,000).

To explore if dQTLs were associated with other changes in the tumor epigenome, we studied histone modifications in primary prostate tumors for H3K27ac (n=92 patients), H3K27me3 (n=76) and H3K4me3 (n=56) and androgen receptor (*AR*; n=88) binding ^44^ (**Supplementary Figure 4a; Supplementary Table 1**). Of the 16 tag dQTLs, 10 overlap active regulatory regions: six dQTLs overlap H3K27ac sites (2-89 patients) of which five also overlap H3K4me3 (1-47 patients) sites (**Supplementary Figure 4c; Supplementary Table 7**). Five dQTLs overlap H3K27me3, one of which overlapped H3K27ac sites in other patients, indicative of bivalent chromatin. We replicated these findings in a second cohort of 48 primary prostate cancer tumors profiled *via* ChIP-Seq for H3K27ac (n=48 patients), H3K4me2 (n=6 patients), H3K4me3 (n=4 patients), FOXA1 (n=10 patients) and HOXB13 (n=9 patients; **Figure 2e**; **Supplementary Figure 4a**; **Supplementary Table 7**). Two of five dQTLs at H3K27ac modification sites demonstrated allelic imbalance specifically in tumor tissue and not in normal tissue, indicative of allele-specific regulation (**Figure 2e**). Further, of the 16 dQTL tag SNPs, 13 overlapped with active regulatory regions and master transcription factor binding sites in four prostate cancer cell lines and one epithelial cell line (**Supplementary Figure 4d**; **Supplementary Table 7**)^45–58^. **Figure 2f** summarizes all dQTLs overlapping transcription factor binding sites or regulatory chromatin, which was similar to that expected by chance (P > 0.40). Thus, a subset of dQTLs may modulate transcription factor binding or histone modifications, known determinants of local somatic mutation rates, but this is not a primary mechanism of their action^59^.

Finally, to begin to elucidate a mechanism of dQTLs we focused on the impact of rs11203152 – associated with loss of *TMPRSS2* – on the local chromatin structure. rs11203152 is in close proximity to multiple chromatin looping sites anchored by RNA Polymerase II (RNAPII), RAD21, AR and ERG in prostate cancer cell lines^38^ (**Figure 2g**). To quantify the observed enrichment of regulatory chromatin loops near rs11203152, we tested if the number of anchors within 1 Mbp of rs11203152 was more than expected by chance (permutation test; n = 100,000 randomly selected regions of equal size). We discovered an enrichment of RAD21 chromatin loop anchors around rs11203152 in LNCaP cells (BH FDR = 0.04; observed number of anchors = 84; expected = 35) but not DU145 cells (BH FDR = 0.19; observed = 66; expected = 28; **Figure 2h**). As LNCaP cells are hormone sensitive prostate cancer cells while DU145 are hormone insensitive, this suggestions rs11203152 may impact AR regulation. VCaP cells, which have a T2E fusion, showed an enrichment of RNA Polymerase II (BH FDR = 0.04; observed = 95; expected = 18), AR (BH FDR = 0.04; observed = 325; expected = 75) and ERG (BH FDR = 0.04; observed = 83; expected = 22) anchored chromatin loops around rs11203152 (**Figure 2h**). These data suggest rs11203152 may interact with AR regulation to promote loss of *TMPRSS2* ^60^.

### dQTLs modulate tumor gene expression

Given the overlap of dQTLs in areas of active chromatin, we sought to quantify their influence on tumor gene expression. We assessed if any dQTL tag SNPs were expression quantitative trait loci (eQTLs) for their associated somatic driver gene (**Supplementary Figure 4e; Supplementary Table 1**). We identified two dQTL-eQTLs associated with *RB1* mRNA abundance and one with *TMPRSS2* (BH FDR < 0.1; **Figure 2f; Supplementary Figures 4f-h**). Both rs12385878 and rs7320595 were associated with RB1 protein abundance (β = 0.29; BH FDR = 7.87×10^−2^; **Supplementary Figure 4i-j**) and rs13048402 was nominally associated with TMPRSS2 protein abundance but did not survive multiple hypothesis testing correction (β = -0.24; BH FDR = 0.11; **Supplementary Figure 4k**). To expand eQTL discovery beyond somatic driver genes we evaluated genes in close proximity to the dQTL, defined as ±500 kbp (**Supplementary Figure 4l**). To our surprise, only a single additional eQTL was significant after correcting for multiple hypothesis testing: rs12653946 – *IRX4* ^61^ (β = -0.79; BH FDR = 7.78×10^−14^; **Supplementary Figure 4m**). To determine if there was broader transcriptome modeling, we quantified dQTL association with distal gene abundances, defined as >500 kbp from the SNP. We identified two distal eQTLs (**Supplementary Figure 4l**): rs11203152 – *COX7B* (β = 0.53; BH FDR = 4.46×10^−2^; **Supplementary Figure 4n**) and rs848047 – *MTRR* (β = 0.38; BH FDR = 4.46×10^−2^; **Supplementary Figure 4o**). **Figure 2f** and **Supplementary Table 7** summarize dQTLs influences on gene-expression. Finally, we leveraged the Genotype-Tissue Expression (GTEx) ^62^ project to evaluate if dQTL tag SNPs were associated with mRNA abundance in non-malignant prostate tissue. Three dQTLs were involved in normal tissue eQTLs, including rs12653946 – *IRX4* (P < 3.8×10^−5^; **Supplementary Table 8**). Thus, a subset of dQTLs may directly modulate the tumor transcriptome and proteome.

We reasoned that if dQTLs provide a fitness advantage, tumors might acquire a similar advantage *via* somatic mutations as well ^63^ **(Supplementary Figure 4a**). To test this hypothesis, we evaluated whether somatic mutations were enriched within the region of individual dQTLs. We focused on the 16 dQTL tag SNPs and identified somatic SNVs within ± 10 kbp of each. The number of somatic SNVs within dQTL elements were compared to the local background mutational burden. While regions harboring dQTLs also harbored multiple somatic SNVs (range: 0-6), we did not observe an enrichment of somatic SNVs above chance (P > 0.15; Poisson generalized linear regression; **Supplementary Figure 5a**). This was consistent in breast (range: 3-18; P > 0.09; **Supplementary Figure 5b**), ovarian (range: 0-19; P > 0.13; **Supplementary Figure 5c**) and pancreatic cancers (range: 2-22; P > 0.06; **Supplementary Figure 5d**). Thus, based on this limited subset of dQTLs, germline dQTLs are not at somatic mutation hotspots.

### dQTL allelic frequencies are biased across ancestry populations

It has been well established that genetic ancestry is associated with specific features of the somatic landscape of prostate cancer ^14–18^, but it is unknown if specific germline SNPs contribute a significant proportion of these differences. We quantified the differences in SNP allele frequencies (VAF) between individuals of European, African and East Asian ancestries for dQTL tag SNPs (**Supplementary Figure 4a; Supplementary Table 7**). All 16 dQTL tag SNPs had significantly different VAF between European and African or East Asian populations (BH FDR < 0.01; Fisher’s exact test; **Supplementary Figure 5e-f**). As a control, only two dQTL tag SNPs, rs439864 and rs7679673, had significantly different VAFs within European populations demonstrating dQTLs are not driven by population stratification (BH FDR = 6.75×10^−3^ and 2.65×10^−2^; **Supplementary Figure 5g**). Leveraging a cohort of 91 men of African descent with localized prostate cancer^31^, we tested the 23 concordant dQTLs. Of these, 13 dQTLs had MAF > 0.05, six showed concordant ORs and none statistically replicated (**Supplementary Figure 5h; Supplementary Table 5**).

We then focused on SNPs associated with two mutations with strong ancestry associations: T2E and *FOXA1* ^14–18^. The T2E gene fusion occurs less frequently in individuals of African and East Asian ancestry. The rs11203152 dQTL was associated with an increased risk of loss of *TMPRSS2* in both discovery and replication cohorts (**Figure 1b & 2a**). Concordant with these ancestry trends, the VAF for this SNP was significantly lower in both African and East Asian populations compared to European (VAF_African_ = 0.066; VAF_East Asian_ = 0.000; VAF_European_ = 0.103; BH FDR < 0.01). We tested the association of rs11203152 with loss of *TMPRSS2* in 115 African men from South Africa, Australia or Brazil with prostate cancer, yielding a near-identical effect-size (OR_African_ = 2.45; P_African_ = 0.13; **Supplementary Figure 5i**). Similarly, *FOXA1* SNVs are more common in men of African ancestry than in men of European ancestry ^17^, while in men of East Asian ancestry a coding hotspot SNV not found in other ancestries is common ^16^. The rs848048 dQTL tag SNP was associated with occurrence of SNVs in the 3’ UTR of *FOXA1* (**Figure 1b & 2a**). Concordant with these ancestry differences, the tag SNP had a significantly lower VAF in African populations than in European or Asian ones (VAF_African_ = 0.231; VAF_European_ = 0.485; VAF_East Asian_ = 0.462; OR = 0.36; BH FDR < 0.1). We tested the association between rs848048 and SNVs in *FOXA1* UTR in 115 African men. The allele distribution was substantially different in African individuals compared to European individuals and the association did not replicate in the African cohort (OR_African_ = 0.96; P_African_ = 1.00; **Supplementary Figure 5j**) supportive of a germline role in ancestry-related somatic differences. Assuming these dQTLs have a similar mechanism across ancestry populations, we estimated that 16.7-31.4% of the ancestral differences in T2E and 0.9-67.3% of the ancestral differences in *FOXA1* may be explained by these individual dQTLs (**Figure 2i**). The low explanatory power of rs848048 in Asian populations may be due to different somatic mutational processes. In men of Asian descent, *FOXA1* mutations were almost exclusively localized to a hotspot immediately after the forkhead domain, whereas mutations spanned the entire gene in individuals of European descent^16^. These data suggest dQTLs offer a way of understanding at least a subset of ancestral differences in cancer landscapes^64^.

### dQTLs are associated with clinical outcome

Given that many somatic mutations and mutational processes are predictive of prostate cancer aggression^37,65^, we evaluated whether dQTLs might predict specific clinical features (**Supplementary Figure 4a**). We first considered biochemical relapse, defined by rising serum PSA levels following primary treatment, which is considered a surrogate for prostate-cancer specific mortality ^66^. One dQTL, rs7320595 associated with clonal loss of *RB1*, was nominally associated with biochemical relapse, but did not survive adjustment for multiple hypothesis testing (HR = 1.52; P = 0.05; Cox PH; **Figure 2f; Supplementary Figure 5k-l**; **Supplementary Table 7**). Four dQTLs associated with subclonal gain of *NCOA2* were associated with ISUP grade group at diagnosis (BH FDR < 3.10 × 10^−2^; **Figure 2f; Supplementary Figure 5m-p**). One dQTL, rs848047 associated with SNVs in the 3’ UTR of *FOXA1*, was nominally associated with risk of prostate cancer diagnosis but did not survive multiple hypothesis testing correction (OR = 1.02; P = 0.05; **Figure 2f; Supplementary Figure 5q**).

### A long tail of dQTLs

dQTL discovery requires matched blood and tumor tissue profiles. Despite using the largest whole-genome sequenced prostate cancer cohort available, the statistical power available is much smaller than modern GWAS cohorts. The low frequency of most prostate cancer somatic drivers (∼5-20%) further reduces the power of our analysis. A cohort of our size would have 80% power to identify local dQTLs with MAF ≥ 0.4 and OR ≥ 1.7 for somatic drivers present in half the population (P < 5×10^−4^; **Supplementary Figure 6a**). For common somatic drivers (5-20% frequency), we have at best 80% power to detect an OR above 2.0 (**Supplementary Figure 6b&c**). We identified 35 dQTLs involving 11 somatic drivers and 27 SNPs (**Figure 2b**). From these figures, we estimate that at least 314 additional dQTLs remain to be discovered in larger cohorts at similar effect-sizes (see **Methods**). Identifying dQTLs genome-wide requires a more stringent p-value threshold (P < 5×10^−8^) than achievable with current cohorts.

Given this large number of potentially undetected dQTLs, we evaluated whether there was evidence for a large landscape of subthreshold candidate dQTLs, as has been seen in many early GWAS analyses. We evaluated the five most recurrent somatic drivers: T2E, clonal loss of *ZNF292*, clonal loss of *RB1*, clonal loss of *NKX3-1* and clonal loss within *TMPRSS2*. For each, we evaluated the distribution of p-values for the linear, spatial and enhancer local dQTL analyses to determine if there were more subthreshold p-values than expected by chance. We compared the skew of the real p-value distributions to empirical null distributions generated by randomly shuffling patient-driver assignments, maintaining driver frequency (**Figure 3a-c; Supplementary Figure 6d-f**). Both T2E and clonal loss of *ZNF292* had significantly skewed p-value distributions (**Figure 3a**). T2E showed a significant skew towards small p-values in linear local dQTL discovery (FC_skew *vs*. null skew_ = 1.41; P = 0.016; **Figure 3b; Supplementary Figure 6g**), while clonal loss of *ZNF292* showed a significant skew towards small p-values in spatial local dQTL discovery (P = 0.046; **Figure 3c; Supplementary Figure 6h**). To supplement, we estimated the proportion of non-null p-values within the p-value distributions for local linear, spatial and enhancer dQTLs for the five most recurrent somatic drivers (*i*.*e*., the proportion of p-values that deviate from a uniform distribution). An estimated 7.0-63.1% of dQTLs tested were estimated to be non-null for clonal loss of *NKX3-1*, clonal loss of *ZNF292*, clonal loss of *TMPRSS2* and T2E (**Figure 3d)**. Non-null p-values for spatial dQTLs associated with clonal loss of *RB1* could not be estimated due to insufficient SNPs tested *via* this strategy. These data suggest many additional prostate dQTLs remain to be identified.

**Figure 3.**
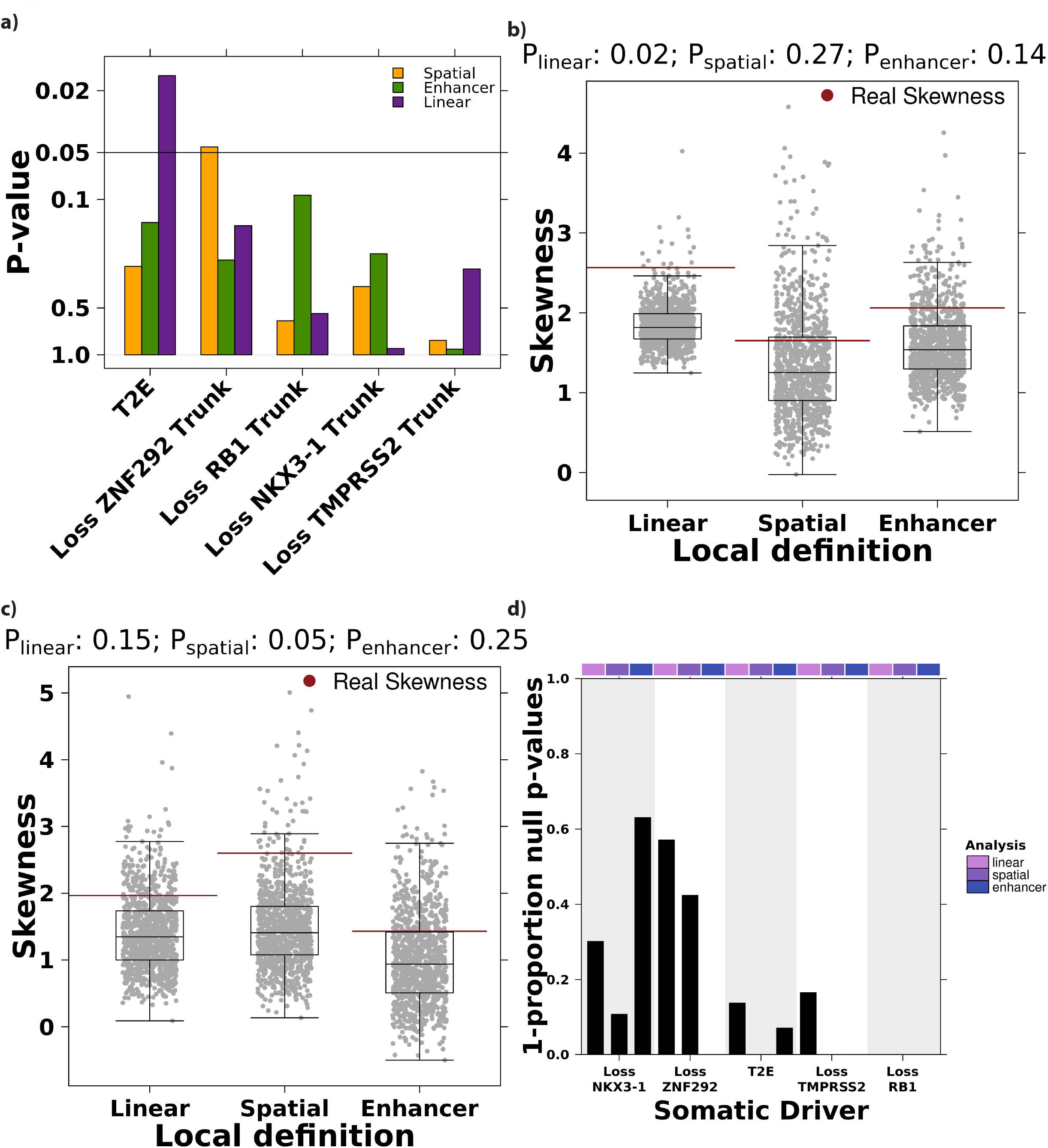
dQTL discovery p-value distribution is significantly skewed. **a)** dQTL discovery p-value distributions are significantly skewed towards smaller p-values. The p-value skew for each dQTL discovery for the top five most recurrent somatic drivers was compared to an empirically generated null distribution (iterations=1,000) and a p-value calculated as the number of null iterations with skew > real skew. Barplot shows the p-value from this permutation analysis. Horizontal line indicates P = 0.05 and colors represent the dQTL discovery approach. **b)** Null skew distribution for T2E dQTL discovery from 1,000 iterations. Horizontal red lines represent real skew values for each dQTL approach. P-values along the top represent the number of null iterations with skew > real skew divided by the number of null iterations. Boxplot represents median, 0.25 and 0.75 quantiles with whiskers at 1.5x interquartile range. **c)** Null skew distribution of clonal loss of *ZNF292*. **d)** Barplot of 1 – the estimated proportion of null p-values (y-axis) in linear, spatial or enhancer dQTL discovery for the top five most recurrent somatic events. The estimated proportion of null p-values of spatial dQTLs associated with clonal loss of *RB1* could not be tested due to too few SNPs.

## Discussion

Every tumor is different, with a life history shaped by its encounters with mutagens, selective microenvironmental pressures, and by stochastic processes^3^. This life history occurs in the context of the patient’s unique germline genome. Subtle differences in germline structure or function have decades to exert their small effects to influence tumor evolution. There are many potential mechanisms of this influence. For example, if an individual possesses germline SNPs that enhance or reduce the efficacy of an oncogenic pathway, cells that acquire a somatic aberration in the same pathway may develop a stronger fitness advantage and experience clonal expansion. Germline SNPs could marginally up- or down-regulate specific pathways making it easier or harder for a tumor genome to further deregulate the same pathway^67^. Similarly, a dQTL deregulating the epigenome or mRNA and protein abundance of a specific gene may change the selective advantage of further amplification or deletion of that gene. These mechanisms are supported by the subset of dQTLs that were additionally meQTLs, eQTLs and pQTLs. Alternatively, if a specific germline SNP renders particular bases of the genome less amenable to DNA damage repair, mutations in these regions may be more likely. dQTLs could affect the structural integrity of the local chromatin, or influence activity of master regulators. This latter example may be illustrated by rs11203152 associated with loss of *TMPRSS2* and located in a region enriched for AR-mediated chromatin looping. This large variety of potential mechanisms supports the idea of polygenic models, where many SNPs modestly influence somatic driver acquisition.

This study focused on identifying limited subsets of driver genes and of dQTLs in prostate cancer and very likely substantially underestimates the full catalogue. First, we only considered somatic drivers present in at least 5% of patients in our discovery cohort. Given the long tail of cancer driver genes^32^, this alone suggests more germline SNPs influence the molecular evolution of prostate cancer. Second, to enrich for promising candidates given limited statistical power, we focused on dQTLs in linear or spatial proximity to driver genes. This allowed us to focus on 5,516 independent genetic SNPs across all 17 somatic drivers: ∼0.6% of the ∼900,000 independent sites tested in a single GWAS^68^. Our statistical efficiency restricted us to finding local dQTLs. Any distal dQTLs remain to be uncovered. Third, our discovery identified dQTLs specifically biasing early *vs*. late mutational events suggesting different germline pressures emerge as a tumor evolves. Limited by available data, we were unable to account for differences in evolutionary timing of drivers during dQTL replication and may have diluted signal of dQTLs as a result. Fourth, the discovery and replication cohorts are comprised of men with low, intermediate and high-risk prostate cancer and the replication cohort included a larger proportion of men with high-risk prostate cancer than the discovery cohort. We are not presently powered to identify dQTLs specific to each risk category and the increased proportion of high-risk prostate cancer in the replication cohort may have reduced the replication of dQTLs specific to low or intermediate-risk prostate cancer. Finally, we focused on germline SNPs present in at least 5% of populations with European ancestry. Our discovery of dQTLs explaining ancestry-differences in somatic mutations motivates further exploration of nature *vs*. nurture by quantifying dQTLs in larger populations of different ethnicities from different geographical localities (*e*.*g*., African nationals *vs*. African Americans).

Cancers arise as stochastic mutagenic processes induce variation within the genome, much but not all of which is successfully repaired. By probabilistically influencing these mutations and the effect of these mutations, germline features help explain why every tumor is different. These data strongly support generation of tumor and normal genomic sequencing of highly diverse genetic populations to quantify how our shared ancestry influences cancer development and further delineate contributions of nature *vs*. nurture.

## Methods

### Discovery patient cohort

The discovery patient cohort was comprised of 427 patients with pathologically confirmed prostate cancer and were hormone naive at time of therapy. All patients underwent image-guided external beam radiotherapy (IGRT) or radical prostatectomy (RadP) with curative intent. The discovery cohort consisted of 276 samples that were published and processed as previously described ^28^, 83 were previously published in Wedge *et al*. ^26^, 50 in Baca *et al*. ^24^, seven in Berger *et al*. ^25^ and 11 in Weischenfeldt *et al*. ^27^. All men were genetically of European descent. Genetic ancestry was determined by calculating genetic distance to well defined populations from the 1000 Genomes Project according to Heinrich *et al*. ^69^. Genetic principal components were determined using plink –pca (v1.9) after pruning for linkage disequilibrium using plink –indep with default parameters.

### Whole-genome sequencing of discovery cohort

For each patient, both blood and tumor sample underwent whole genome sequencing as previously described ^28^. FASTQ files were retrieved for each sample and processed consistently. Raw sequencing reads were aligned to the human reference genome, hs37d5, using BWA-mem ^70^ (v0.7.12-0.7.15) at the lane level. Lane level bam files were merged across libraries with duplicates marked within libraries using Picard (v1.121-2.8.2). Local realignment and base quality recalibration were completed on tumor/normal pairs together with the Genome Analysis Toolkit ^71^ (GATK v3.4.0-3.7.0). Tumor and normal samples were extracted separately, headers corrected (SAMtools v0.1.9-1.5) ^72^ and files indexed (Picard v2.17.11) into individual sample-level BAMs. Finally, sequencing coverage was computed using Picard (v2.17.11) CollectRawWgsMetrics with the default cut-off.

### Germline SNP detection in discovery cohort

Germline SNPs were first identified using GATK (v3.4.0-3.7.0) for each patient individually using HaplotypeCaller followed by VariantRecalibration and ApplyRecalibration ^71^. Individual VCFs were merged using BCFtools (v.1.8) assuming SNPs not present in an individual VCF were homozygous reference. The minor allele frequency (MAF) in the discovery cohort of all SNPs within the merged VCF was calculated and filtered to consider only SNPs with MAF > 0.01 based on the discovery cohort (n=10,058,344). Next, all patients were re-genotyped using GATK (v.4.0.2.1) at these sites to produce gVCFs (*i*.*e*., with option -ERC GVCF). Individual gVCFs were merged using GenomicsDBImport and joint genotyping was run using GenotypeGVCFs. Finally, SNPs were recalibrated using VariantRecalibrator and ApplyVQSR. We determined pathogenic variants within National Comprehensive Cancer Network (NCCN) prostate cancer predisposition genes based on “pathogenic” or “likely pathogenic” annotations in ClinVar and ensuring more than one submitter (*i*.*e*., review status ≥ 2/4 stars).

### Somatic variant detection in discovery cohort

Somatic variants were detected as previously described ^28^. Briefly, somatic single nucleotide variants (SNVs) were detected with SomaticSniper (v1.0.5) with mapping quality threshold set to one and with all other parameters set to their defaults ^73^. SNVs were filtered using LOH, read count and high confidence filters provided with the SomaticSniper package. SNVs were further filtered using in-house filters to account for read coverage, germline contamination, mappability, among others. A full description of these filters can be found here ^28^. Small Indels were identified with cgpPindel v2.2.4 ^74^ with default parameters and the following genomic rules (F004, F005, F006, F010, F012, F018, F015, F016) and soft results (F017). Indels were filtered based on a panel of non-cancer reference samples (pindel_np.gff3.gz), simple repeats, band anchors and germline contamination, amongst others. A full description of these filters can be found here ^28^. SNVs and Indels were annotated to genes using SnpEff (v4.3R) ^75^. Somatic copy number alterations (CNAs) were identified using Battenberg (cgpBattenberg v3.3.0, BattenBerg R-core v2.2.8, alleleCount v4.0.1, PCAP-core v4.3.2, cgpVcf v2.2.1, impute2 v2.3.3) ^76^. Clonal (*i*.*e*., trunk) and subclonal (*i*.*e*., branch) CNAs were predicted using the default cut-off of p-value 0.05 and segments length below 10kb were filtered out.

### Recurrent somatic drivers in prostate cancer

We considered a set of 180 somatic drivers were identified in 666 localized prostate tumors ^28^, and included those with a frequency ≥ 5% in the discovery cohort that has been previously reported in localized prostate cancer (15-19,26-30,33,40,46). Representative genes were selected for CNA drivers based on recurrent regions prioritized by GISTIC and previous literature on the CNA landscape of localized prostate cancer. CNAs represented by genes may be arm level chromosome alterations, such as loss of NKX3-1 which often represents loss of the p-arm of chromosome 8 ^31,40^. This resulted in analysis of 17 somatic drivers: 11 CNA losses (seven trunk and four branch), three CNA gains (two trunk and one branch), one fusion (the recurrent T2E fusion between *TMPRSS2* and *ERG*) and two SNVs. For a full definition of each somatic driver refer to **Supplementary Table 1**.

### dQTL discovery: risk SNPs dQTLs

The 147 SNP polygenic risk score generated by Schumacher *et al*. ^9^ was first considered for dQTL discovery. Of the 147 SNPs, 135 had a MAF > 0.05 in the discovery cohort. All 135 SNPs were tested for association with all 17 somatic drivers using a logistic regression model correcting for the first five genetic principal components, age and mutation burden. P-values were adjusted for multiple-hypothesis testing using Benjamini & Hochberg false discovery correction. Significance was defined as BH FDR < 0.1.

### dQTL discovery: linear local dQTLs

Local dQTLs were first defined based on the linear orientation of the genome. Considering each somatic event could be defined by a single gene, germline SNPs within ±500 kbp of the affected gene were interrogated for their association with the somatic event. Associations were quantified using a logistic regression model correcting for the first five genetic principal components, age and the somatic mutation burden (*i*.*e*., PGA when testing CNAs and SNV mutation density when testing SNV). Haplotype blocks within the defined linear local region were calculated considering the definition by Gabriel *et al*. ^77^ and a Bonferroni threshold considering α = 0.1 was used to determine significance for each somatic driver. We selected α=0.1 as our significance threshold to reduce false negatives in our discovery given the relatively small size of our cohort. All discovered dQTLs were tested in an independent replication cohort in order to remove likely false positives. Discovery dQTL statistics for all tested SNPs – unpruned for linkage disequilibrium – are provided in **Supplementary Table 4**.

### dQTL discovery: spatial local dQTLs

Local dQTLs were defined taking into consideration the three-dimensional structure of DNA. The term spatial local was defined as regions of the DNA, outside ±500kbp around the affected gene, that loop to interact with the driver gene. First, these regions were defined by RAD21 ChIA-PET profiling in LNCaP and DU145 cell lines ^39^ and RNA polymerase II ChIA-PET profiling in LNCaP, DU145, VCaP and RWPE-1 cell lines ^38^. Coordinates of driver genes were overlapped with peak anchor regions using BEDtools. Based on an interaction map, peak anchors paired with driver-gene-overlapped peaks were defined as interacting regions. Similar to linear local dQTLs, associations were quantified using a logistic regression model correcting for the first five genetic principal components, age and the somatic mutation burden. Again, haplotype blocks within the defined spatial local region were calculated considering the definition by Gabriel *et al*. ^77^ and a Bonferroni threshold considering α=0.1 was used to determine significance for each somatic driver. Discovery dQTL statistics for all tested SNPs – unpruned for linkage disequilibrium – are provided in **Supplementary Table 4**.

### dQTL discovery: enhancer local dQTLs

Spatial local regions were defined based on HiChIP H3K27ac profiling in LNCaP cell lines. HiChIP was conducted as reported previously. Again, associations were quantified using a logistic regression model correcting for the first five genetic principal components, age and the somatic mutation burden and haplotype blocks within the defined enhancer local region were calculated considering the definition by Gabriel *et al*. ^77^ and a Bonferroni threshold considering α = 0.1 was used to determine significance for each somatic driver. Discovery dQTL statistics for all tested SNPs – unpruned for linkage disequilibrium – are provided in **Supplementary Table 4**.

### Prostate cancer replication cohort

Individuals of European descent, as determined by Yuan *et al*. ^64^, from TCGA PRAD project were used as a replication cohort ^31^. As described previously ^23^, concordance between SNP6 microarray (SNP6) genotypes and whole exome sequencing (WXS) of blood sample genotypes was evaluated and only samples with >80% concordance were retained (n = 412 samples). Genotypes, from SNP6 supplemented by WXS, were imputed using the Michigan Imputation Server – pre-phasing using Eagle (v2.4) ^78^, imputation using Minimac4 ^79^ and the Haplotype Reference Consortium (release 1.1) panel ^80^. A final list of 40,401,582 SNPs was then available for validation studies. SNV and CNA calls based on hg19 reference genome were downloaded from GDC Archive legacy (https://portal.gdc.cancer.gov/legacy-archive/search/). T2E fusions for TCGA samples were identified using FusionCatcher (v.0.99.7c) ^81^. A second cohort of 140 Australian men with localized prostate cancer was used to supplement the replication cohort. All men were of European descent as determined according to Heinrich *et al*. ^69^. All patients had blood and tumor WGS that was processed with the same pipelines as the discovery cohort, including evolutionary timing of CNAs ^28^. Similar to the discovery cohort, germline SNPs were identified using GATK (v3.4.0-3.7.0) ^71^. First, HaplotypeCaller was run on the normal and tumor BAMs together, followed by Variant Recalibration and ApplyRecalibration, following GATK best practices. Germline SNPs were filtered for somatic and ambiguous variants that had more than one alternate base. Genetic principal components were determined using plink –pca (v1.9) after pruning for linkage disequilibrium using plink –indep with default parameters.

### Pan-cancer replication cohort

We leveraged the Pan-Cancer Analysis of Whole Genomes (PCAWG) ^32^ to test the replication of dQTLs in other cancer types, using germline VCFs and somatic CNA calls from the Pan-Cancer Analysis of Whole Genomes from DCC (https://dcc.icgc.org/releases/PCAWG/). We considered only adult cancers with >100 samples: breast, ovarian, pancreatic and liver cancer. Next, we only considered patients of European ancestry which resulted in 134 breast, 91 ovarian, 116 pancreatic and 0 liver cancer patients. Thus, we did not consider liver cancer in replication analysis. We tested somatic events with a recurrence rate ≥ 5% in each cancer type.

### Replication of dQTLs

dQTLs with available somatic profiling and germline genotyping were tested in the replication cohort. TCGA does not have WGS so the evolutionary timing of CNAs could not be determined in these patients. Thus, dQTLs involving CNAs were tested in TCGA without considering trunk *vs*. branch classifications. As a result, there were significant differences in the proportion of cases and controls between the discovery and replication cohorts (**Supplementary Table 1**). T2E calls for TCGA samples in the replication cohort were based on RNA-sequencing alone compared to the rest of the samples which considered DNA sequencing or the union of DNA and RNA sequencing when available. dQTLs in all replication cohorts were tested using the same logistic regression model as used in discovery, correcting for the first five genetic principal components, age and the total burden of somatic mutation type being tested (*i*.*e*., PGA or SNV mutation density). dQTLs were considered to have replicated if BH FDR < 0.1 and sign(log(OR_discovery_)) = sign(log(OR_replication_)).

### Replication of dQTLs in ICGC EOPC

We identified nine dQTLs that were associated with somatic events with a recurrence rate ≥ 5% in the EOPC-DE cohort and had concordant ORs in the discovery and replication cohorts. The candidate SNPs were studied across 238 prostate cancer patients with European ancestry from the ICGC EOPC-DE cohort ^41^. Germline SNP genotyping and quality control was performed as previously described ^82^. Association between germline SNP genotypes and presence of somatic mutation was performed using logistic regression models in Python (stats package version 0.11.1) correcting for the first five principal components, age and mutational burden.

### Replication of dQTLs in Hartwig Medical Foundation metastatic prostate cancer

We replicated dQTLs on the external CPCT-02/HMF dataset under data-requests DR-071 and DR-208 ^42^. This as an extension of the metastatic prostate cancer cohort (*n* = 394 distinct patients) as previously described by van Dessel & van Riet *et al* ^83^. To select patients of (predominantly) European descent, we utilized the established set of ancestry markers from the EUROFORGEN Global AIM-SNP set ^84^ which consisted out of 128 bi-allelic and tri-allelic germline markers and 934 respective reference samples of African, East Asian, European, Native American and Oceanian ancestry. For these ancestry markers, we determined the respective germline genotype (0/0, 0/1, 1/1, 0/2, 1/2 or 2/2) within all distinct patients within the CPCT-02/HMF dataset. Subsequently, we performed a Principal Component Analysis (PCA) on the combined dataset of genotypes from the CPCT-02/HMF dataset and reference samples. As input for the PCA, genotypes were converted into six numerical categories (0 to 5) and zero centered and scaled during PCA. To determine the putative ethnicity of the CPCT-02/HMF patients, we performed a K-Means clustering (*k* = 5, Hartigan and Wong algorithm on 50 random sets and 10,000 iterations) on all principal components (*i*.*e*., ancestry markers) as derived on the combined genotype-matrix of the reference samples and the CPCT-02/HMF dataset. From this analysis, we selected the distinct CPCT-02/HMF patients clustering within the European descent reference-cluster (*n* = 384). For these 384 European CPCT-02/HMF metastatic patients, we determined the germline genotypes of the dQTLs (*n* = 19) and the presence of the respective somatic event within the tumor genome (somatic deletions of *CDKN1B, CHD1, RB1, TMPRSS2* and/or *ZNF292*, amplifications of *NCOA2*, somatic mutations within the 3’ UTR of *FOXA1* and genomic fusions of *TMPRSS2*-*ERG*). If multiple metastatic biopsies from the same patients were available (*n* = 43), the aggregation of respective somatic events within a patient was used to determine the presence of these somatic events. dQTLs were assessed within a logistic regression model correcting for the first five genetic principal components (based on the ancestry markers), age and mutational burden.

### Replication of dQTLs in PROFILE

Dana-Farber Cancer Institute prospective cohort (PROFILE) was collected with informed consent: 490 unrelated men of European descent with prostate cancer (91 with metastatic disease and 399 primary or local tumors). All samples underwent targeted sequencing on the OncoPanel platform with three panel versions that targeted the exons of 275, 300 and 447 genes, respectively. Genotypes were imputed from off-target reads using STITCH (v.1.5.3) ^85^. To determine genetic ancestry, reference principal components were computed by SNPweight tools in HapMap populations of European, West African (Yoruban) and East Asian (Chinese) ancestry ^86^. In the PROFILE cohort, imputed dosages for variants with INFO > 0.4 and MAF > 0.01 were projected the same PCA space using the PLINK2 ‘--score’ function. The mean principal component along both the West African-European cline and the East-Asian-European cline was computed for all individuals who self-reported as white. Individuals within ± two standard deviations were retained. Samples were filtered for relatedness using a GRM matrix with a 0.05 cutoff. SNPs were filtered to ensure Hardy-Weinberg equilibrium p-value > 0.001, MAF > 0.05 and INFO > 0.4. If the tag dQTL was not genotyped in the PROFILE cohort, a proxy SNP was selected by maximizing the product of the INFO, R^2^ and 1000 Genomes European MAF using LDlinkR ^87^. Finally, associations were tested using a logistic regression in PLINK2 with the first five genetic principal components, tumor purity, panel version, age and PGA as covariates.

### Meta-analysis across discovery, replication, HMF, EOPC and PROFILE cohorts

Effect sizes and standard errors of dQTL associations in the discovery, replication, HMF, EOPC and PROFILE cohorts were combined using a restricted maximum likelihood model as implemented in the metafor R package (v3.0.2).

### Chromothripsis associations

We tested the association of dQTLs with chromothripsis as determined in Fraser *et al*. ^40^. We applied a linear regression model correcting for the first five principal components and age.

### Germline methylation (meQTL) associations

To assess the effect of dQTLs on the tumor methylome, the 16 concordant tag dQTLs were evaluated for local meQTLs, defined as probes ±500 kbp around the SNP, using a linear regression correcting for the first five genetic principal components and age. P-values were adjusted for multiple-hypothesis testing using Benjamini & Hochberg (BH) false discovery correction. Significance was defined as BH FDR < 0.10. Significant meQTLs were next replicated in the TCGA cohort using the same linear regression modeling. Here replication was defined as BH FDR_replication_ < 0.10 and sign(β_replication_) = sign(β_discovery_). Replicated meQTLs were tested for tumor specificity considering patients that had matched tumor/reference methylation profiling (n=50). Tumor specificity was defined as BH FDR_tumor_ < 0.10 and BH FDR_reference_ > 0.10 or sign(β_tumor_) ≠ sign(β_reference_) using the same linear regression model. To assess enrichment of meQTLs, we generated a null distribution of the number of SNPs involved in a replicated meQTL and the number of replicated meQTLs. We randomly sampled 16 SNPs from the total list of SNPs evaluated as a local dQTL against any driver. We identified and replicated local meQTL ±500 kbp around each of the 16 random SNPs using the same methods as the dQTL-meQTL analysis. We calculated the number of SNPs involved in a replicated meQTL as well as the total number of replicated meQTLs. We repeated this 1,000 times. P-values were calculated as 1 – the fraction of iterations more dQTLs were involved in a replicated meQTL than random SNPs or 1-the fraction of iterations dQTLs were involved in more replicated meQTLs than random SNPs.

### Germline-chromatin associations

Peak BED files for H3K27ac (n = 92), H3K27me3 (n=76), AR (n=88) and H3K4me3 (n=56) were downloaded for an independent cohort of 94 localized prostate cancer patients from the Gene Expression Omnibus (GSE120738) ^44^. dQTLs overlapping each target were identified using the downloaded bed files. We considered a dQTL overlapping if any of the SNPs in its haplotype block overlapped the target. A second cohort of 48 localized prostate cancer patients was additionally profiled, as described previously ^23^. Briefly, both adenocarcinoma and non-malignant prostate tissue from each patient was subjected to ChIP-Seq for H3k27ac (n=48), H3k4me2 (n=6), H3k4me3 (n=4), FOXA1 (n=10) and HOXB13 (n=9) and blood samples were genotyped for germline SNPs followed by imputation using the HRC panel ^80^. Sites of allelic imbalance in the ChIP-Seq peaks were identified by first correcting for mapping bias using the WASP pipeline ^88^, peak calling using MACS2 and finally testing for allele-specific signal using GATK ASEReadCounter ^71^ and a beta-binomial test. Each test was performed once for samples from normal, tumor, or both, as well as a test for difference in imbalance between tumor and normal. Peaks were considered “imbalanced” in each of these four test categories if any of the SNPs tested for that peak exhibited allele-specific signal at a 5% BH FDR. Finally, we tested the overlap of dQTLs with published ChIP-Seq data from LNCaP, PC3, 22Rv1, VCaP and RWPE-1 cell lines ^45–58^. If multiple target:treatment pairs existed the median number of overlapping SNPs was used. For all ChIP-Seq analyses, dQTLs were considered overlapping if any of the SNPs within the entire LD block overlapped with the ChIP-Seq peak.

### Germline-RNA (eQTL) and germline-protein (pQTL) associations

Next, the 16 SNPs involved in the 23 concordant dQTLs were tested for their effect on the transcriptome. We evaluated local eQTLs, defined as genes ±500 kbp around the SNP. mRNA abundance TPM values for each gene were rank inverse normalized. eQTLs were tested using a linear regression model correcting for the first five genetic principal components, age and ten PEER ^89^ factors to adjust for noise in the RNA-Seq data. P-values were adjusted for multiple-hypothesis testing using the Benjamini & Hochberg false discovery correction. Nominally significant eQTLs were considered for pQTL discovery using protein abundances from mass spectrometry as described previously ^90^. pQTLs were tested using a linear regression model correcting for the first five genetic principal components, age and ten PEER factors to adjust for noise in the mass spectrometry data.

### Germline-clinical associations

Germline SNPs in dQTLs were associated with clinical characteristics including PSA, ISUP grade group, T-category, age at diagnosis and biochemical recurrence. PSA and age were tested using a linear regression model, correcting for the first five genetic principal components. The PSA model was also corrected for age. T-category was tested using a logistic regression model comparing T2 to ≥T3, correcting for the first five genetic principal components and age. ISUP was tested by using an ordinal linear regression model, correcting for the first five genetic principal components and age. Each clinical outcome was independently corrected for multiple hypothesis testing using the Benjamini & Hochberg false discovery correction. Survival analysis with biochemical recurrence was tested using a Cox Proportional Hazards model. Three genetic models, dominant, recessive and co-dominant, were tested and the model with the lowest AIC was reported. Kaplan-Meier curves were plotted, and HR adjusted for primary treatment.

### Somatic SNV enrichment

For each of the 16 SNPs involved in the high confidence dQTLs, we assessed if the somatic SNV mutation burden ±10 Mbp of the dQTL was higher than expected. We leveraged ActiveDriverWGS ^91^ which uses a Poisson regression to compare the mutation burden of a region of interest to the adjacent genomic window (± 50 kbp). The narrow adjacent window reflects similar chromatin, structure and replication timing to the region of interest. ActiveDriverWGS also corrects for differences in the trinucleotide contexts of the region of interest compared to the flanking windows. P-values were adjusted for multiple hypothesis testing using Benjamini & Hochberg false discovery correction.

### Ancestral variant allele frequency bias

Variant allele frequencies in European (n=7,718), African (n=4,359) and East Asian (n=780) populations for the 16 dQTL SNPs were extracted from gnomAD (v2.1.1) ^92^. Allele frequencies in African and East Asian populations were compared to European population using Fisher’s exact test and BH FDR was applied across all 16 SNPs in each comparison separately. As a control, North-West European VAFs were compared again Other Non-Finnish European VAFs using Fisher’s exact test. These two European populations were chosen because they had the largest sample number in gnomAD. To estimate the proportion of ancestral differences in T2E and *FOXA1* mutation frequency explained by dQTLs, we compared the ORs of ancestry-somatic associations and dQTLs ORs multiplied by normalized variant allele frequency differences between the two ancestry groups. For example:

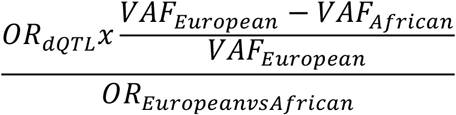

We estimated OR_European *vs*. African_ (T2E) = 5.00 and OR_European *vs*. African_ (*FOXA1* SNVs) = 0.50 based on Huang *et al*. ^15^ and Lindquist *et al*. ^17^ compared to the somatic driver frequency in the discovery cohort. We estimated OR_European *vs*. East Asian_ (T2E) = 7.47 and OR_European *vs*. East Asian_ (*FOXA1* SNVs) = 0.07 based on Li *et al*. ^16^ compared to the somatic driver frequency in the discovery cohort.

### dQTL power analysis

Power was estimated based on the non-centrality parameter of the χ^2^ statistic under the alternative hypothesis using the R package gwas-power (https://github.com/kaustubhad/gwas-power). Power was calculated for varying MAF and effect size values considering sample sizes reflective of somatic driver frequencies 0.05, 0.20 and 0.50 in the discovery cohort. To estimate the number of non-detected dQTLs, discovered dQTLs were binned based on their MAF, effect size and somatic driver frequency and the number of detected dQTLs in each bin was divided by the corresponding power to estimate the total number of dQTLs expected. Next, we subtracted the number of discovered dQTLs from the total number of dQTLs to estimate the number of non-detected dQTLs.

### Assessment of skew of dQTL p-value distributions

To determine if dQTL p-value distributions were significantly skewed to small p-values more than expected by chance alone, a null distribution for each analysis (*i*.*e*., linear local and spatial local) and each somatic driver was generated by permuting the somatic driver labels. That is, for a single somatic event, patients were randomly assigned whether or not they had the somatic event while maintaining the true frequency of the event in the cohort. Next, both linear and spatial local dQTL discovery was conducted as described above with the permuted somatic driver labels. The skew of the -log_10_ p-value distribution was calculated and compared to the true distribution. P-values were calculated by considering the number of permutation iterations that had skew > real skew divided by the number of iterations performed. One thousand iterations were performed for each somatic driver. To supplement these analyses, we also estimated the proportion of null p-values in the p-value distributions for linear, spatial and enhancer dQTLs for the top five most recurrent somatic mutations using the pi0est() function in the qvalue R package (v2.18.0).

## Data visualization

Visualizations were generated in the R statistical environment (v3.3.1) with the lattice (v0.24-30), latticeExtra (v0.6-28) and BPG (v5.6.23) packages ^93^.

## Data Availability

Raw sequencing data are available in the European Genome-phenome Archive under accession EGAS00001000900 (https://www.ebi.ac.uk/ega/studies/EGAS00001000900). Processed variant calls are available through the ICGC Data Portal under the project PRAD-CA (https://dcc.icgc.org/projects/PRAD-CA). Methylation data are available in the Gene Expression Omnibus under accession GSE84043. TCGA WGS/WXS data are available at Genomic Data Commons Data Portal (https://gdc-portal.nci.nih.gov/projects/TCGA-PRAD). Primary samples ChIP-Seq data was retrieved from Gene Expression Omnibus under accession GSE120738.

## Authors’ contributions

**Project Initiation:** K.E.H., P.C.B

**Sample Preparation:** N.K., A.S., S.G.R., J.S., A.J.C., M.M.P, S.B.A.M., P.D.S., R.M.S.B.

**Bioinformatic Analyses:** K.E.H., J.Y., T.S., J.L., N.S.F., J.v.R., K.T., J.S.P., C.H.J., H.Z., T.N.Y., L.E.H., R.J., C.B., E.O., W.J., J.J., S.M.W.

**Statistical Analyses:** K.E.H., J.Y., J.W., J.v.R., K.T.

**Manuscript First Draft:** K.E.H.

**Supervised Research:** B.J.P., N.Z., A.G., M.P.L., A.U.K., N.M.C., R.G.B., S.M.W., J.W., R.H., H.H.H., V.M.H., B.P., M.L.F., C.M.H., R.S.M., P.C.B.

**Manuscript Editing & Approval:** All authors

## Acknowledgements

The authors thank Anamay Shetty and all members of the Boutros lab for helpful suggestions and support. The results described here are based in part upon data generated by the TCGA Research Network: http://cancergenome.nih.gov/. This work was supported by Prostate Cancer Canada and is proudly funded by the Movember Foundation (grant #RS2014-01 to P.C.B.). P.C.B. was supported by a Terry Fox Research Institute New Investigator Award and a CIHR New Investigator Award. This project was supported by Genome Canada through a Large-Scale Applied Project contract to P.C.B., R. Morin and S. P. Shah. K.E.H was supported by a CIHR Vanier Fellowship. This work was supported by the National Health and Medical Research Council (NHMRC) of Australia, grants #APP1165762 to V.M.H. and #APP1104010 and #APP1162514 to C.M.H., and Cancer Association of South Africa (CANSA) to V.M.H. This work was supported by the NIH/NCI under award number P30CA016042, by grants from the National Cancer Institute EDRN (U01CA2141941) and ITCR (U24CA248265). H.H.H. holds Joey and Toby Tanenbaum Brazilian Ball Chair in Prostate Cancer. This work is supported by a Terry Fox New Frontiers Program Project Grant (1090 P3 to H.H.H.). This work was supported by a Prostate Cancer Foundation Special Challenge Award to PCB (Award ID #: 20CHAS01) made possible by the generosity of Mr. Larry Ruvo. This publication and the underlying study have been made possible partly on the basis of the data the Hartwig Medical Foundation and the Center of Personalized Cancer Treatment (CPCT) have made available to the study. N.Z. and K.T. were supported by NIH grants U01HG009080, R01HG006399, R01CA227237, R01ES029929, R01CA227466, R01HG011345 and the DoD grant W81XWH-16-2-0018. This work was supported by W81XWH-19-1-0565, R01CA193910, R01CA251555 and H.L. Snyder Medical Research Foundation to M.L.F. B.J.P. was supported by a Victorian Health and Medical Research Fellowship. S.M.W. was supported by the Research Council of Norway (187615), the South-Eastern Norway Regional Health Authority, and the University of Oslo. R.S.M. acknowledges funding support from NCI grant (R01CA245294), CPRIT Individual Investigator Research Award (RP190454), US Department of Defense Impact Award (W81XWH-17-1-0675) and US Department of Defense Breakthrough Award (W81XWH-21-1-0114).

