## SupplementaryMaterials for "Germline determinants of the prostate tumor genome"

### Supplementary Figure 1

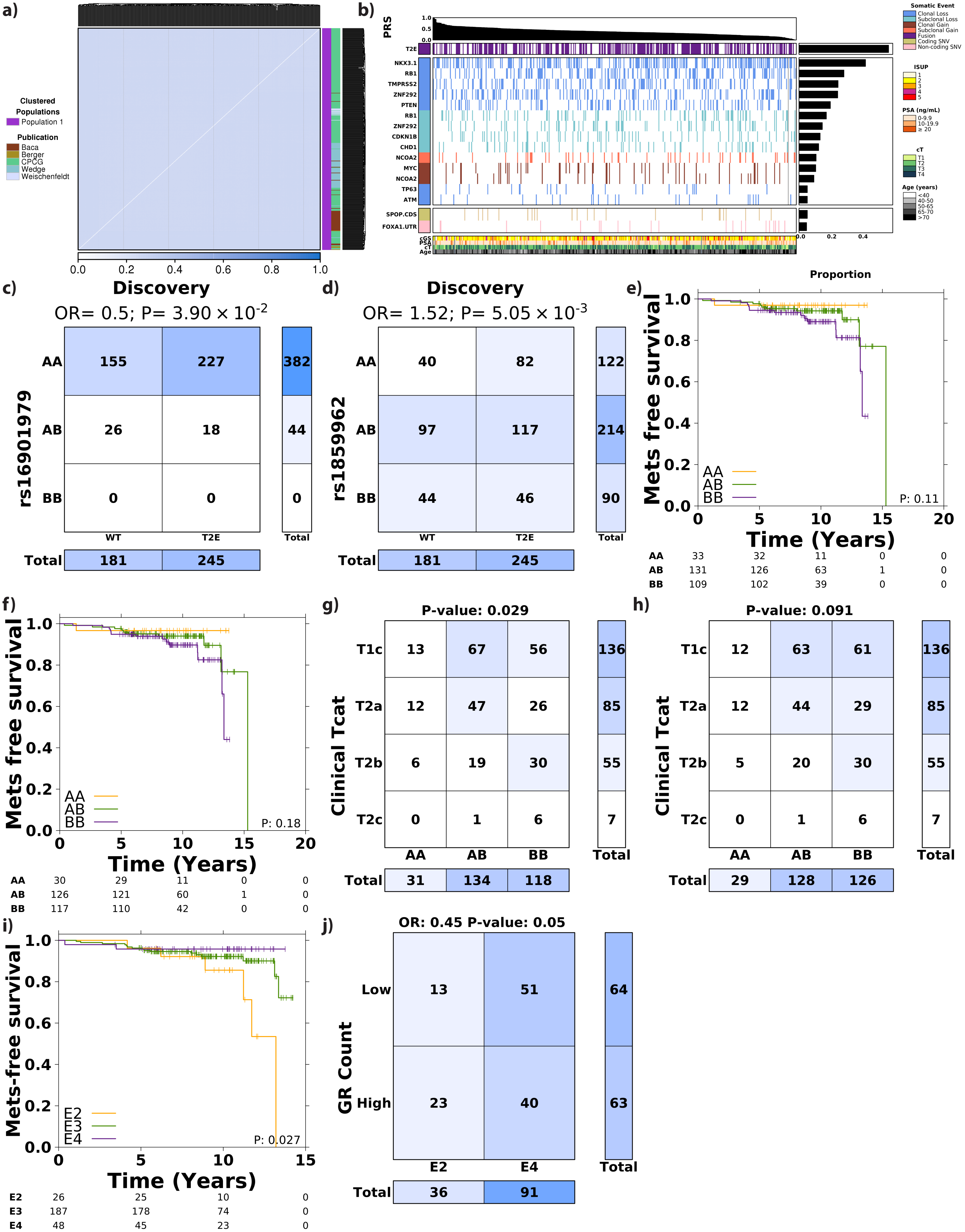

**Supplementary Figure 1 – Cohort characteristics and risk dQTL replication**
**a)** Clustering using identity-by-state (IBS) as the distance metric showed no evidence of population substructure. Heatmap shows the identity-by-state values for all pairwise comparisons. The first covariate along the right shows the cluster provided by plink (v1.9). The second covariate indicates the original cohort the patient was published in. **b)** Landscape of somatic drivers in the discovery cohort. Somatic drivers are categorized as losses (blue), gains (red), SVs (purple), non-coding SNVs (pink) or coding SNVs (khaki). Barplot on the right shows the frequency of each driver in the discovery cohort. Covariate on the bottom indicates clinical characteristics of each patient including clinical ISUP grade group (ISUP), pre-treatment prostate serum antigen (PSA), clinical T category (cT) and age. Barplot on the top indicates the polygenic risk score (PRS), scaled between 0-1, for each patient. **c-d)** Contingency tables of rs16901979 **(c)** and rs1859962 **(d)** associated with T2E in discovery cohort. **e-f)** Kaplan-Meier plots of rs1856888 **(e)** and rs1047303 **(f)** associated with metastasis-free survival (MFS). P-value from log-rank test. **g-h)** Contingency tables of rs1856888 **(g)** and rs1047303 **(h)** associated with clinical T category (Tcat). P-values from Fisher's exact test. **i)** Kaplan-Meier plot of APOE genotypes associated with metastasis-free survival. P-value from log-rank test. **j)** Contingency table of APOE2 and APOE4 associated with GR count. OR and p-value from Fisher's exact test.

**Supplementary Figure 2**

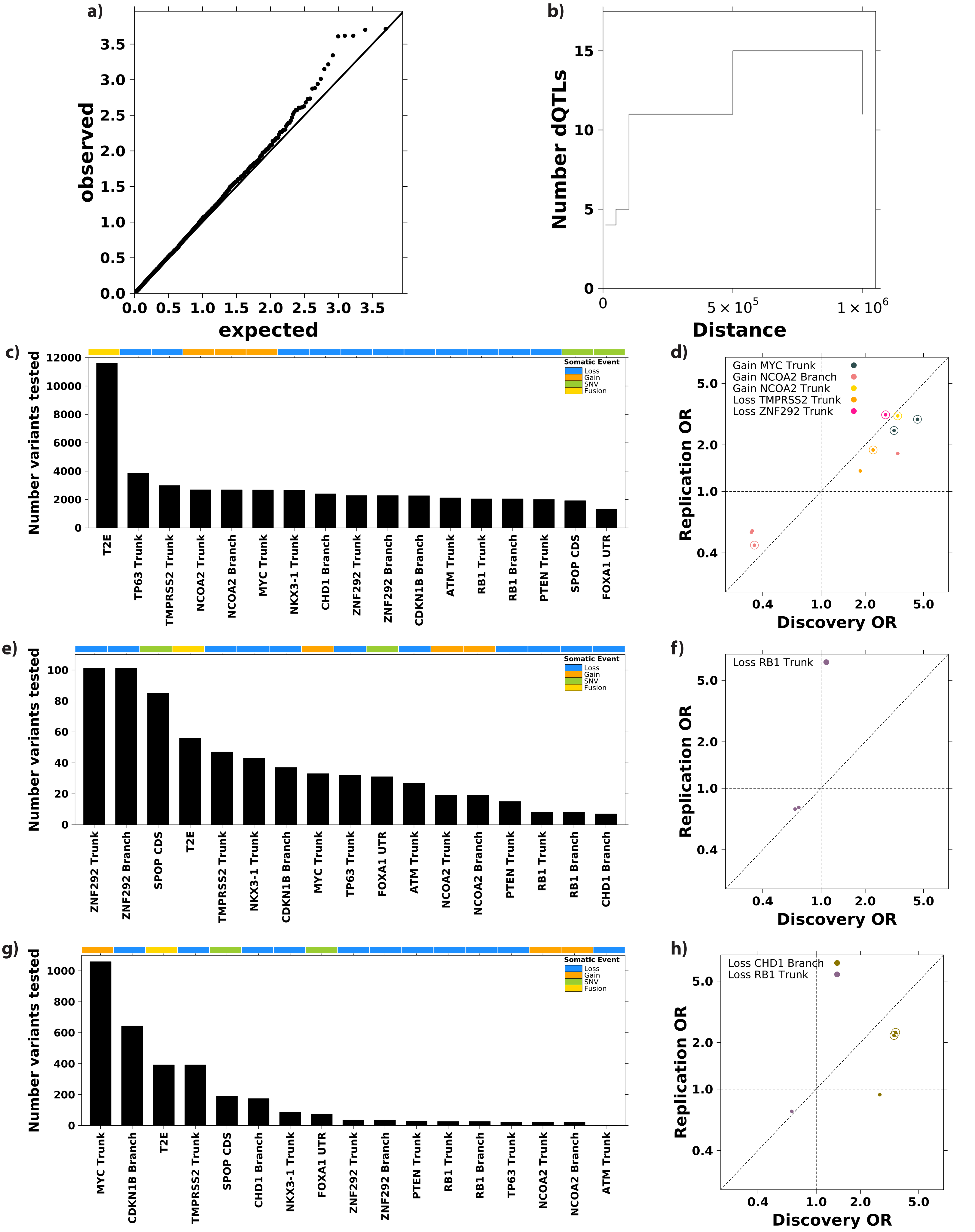

**Supplementary Figure 2 – Local dQTLs discovery**

**a)** QQ plot of expected $-\log_{10}$ p-values *vs*. observed $-\log_{10}$ p-values for association of individual PRS SNPs with 17 drivers. **b)** Sensitivity plot showing number of discovered tag linear local dQTLs based on increasing distance from gene boundaries. **c)** Barplot shows number of variants tested per somatic driver based on linear definition of local dQTL. Covariate along the top indicates the type of somatic driver event. **d)** Comparison of ORs for linear local dQTLs with CNA drivers based on WGS profiling, x-axis and array profiling, y-axis. Horizontal and vertical dotted lines represent OR = 1 and diagonal line represents y=x. Halo around points indicates BH FDR < 0.1 in array-profiled cohort. **e)** Barplot shows number of variants tested per somatic driver based on spatial definition of local dQTL. Covariate along the top indicates the type of somatic driver. **f)** Comparison of ORs for spatial local dQTLs with CNA drivers based on WGS profiling, x-axis and array profiling, y-axis. Horizontal and vertical dotted lines represent OR = 1 and diagonal line represents y=x. Halo around points indicates BH FDR < 0.1 in array-profiled cohort. **g)** Barplot shows number of variants tested per somatic driver based on enhancer definition of local dQTL. Covariate along the top indicates the type of somatic driver. **h)** Comparison of ORs for enhancer local dQTLs with CNA drivers based on WGS profiling, x-axis and array profiling, y-axis. Horizontal and vertical dotted lines represent OR = 1 and diagonal line represents y=x.

### Supplementary Figure 3

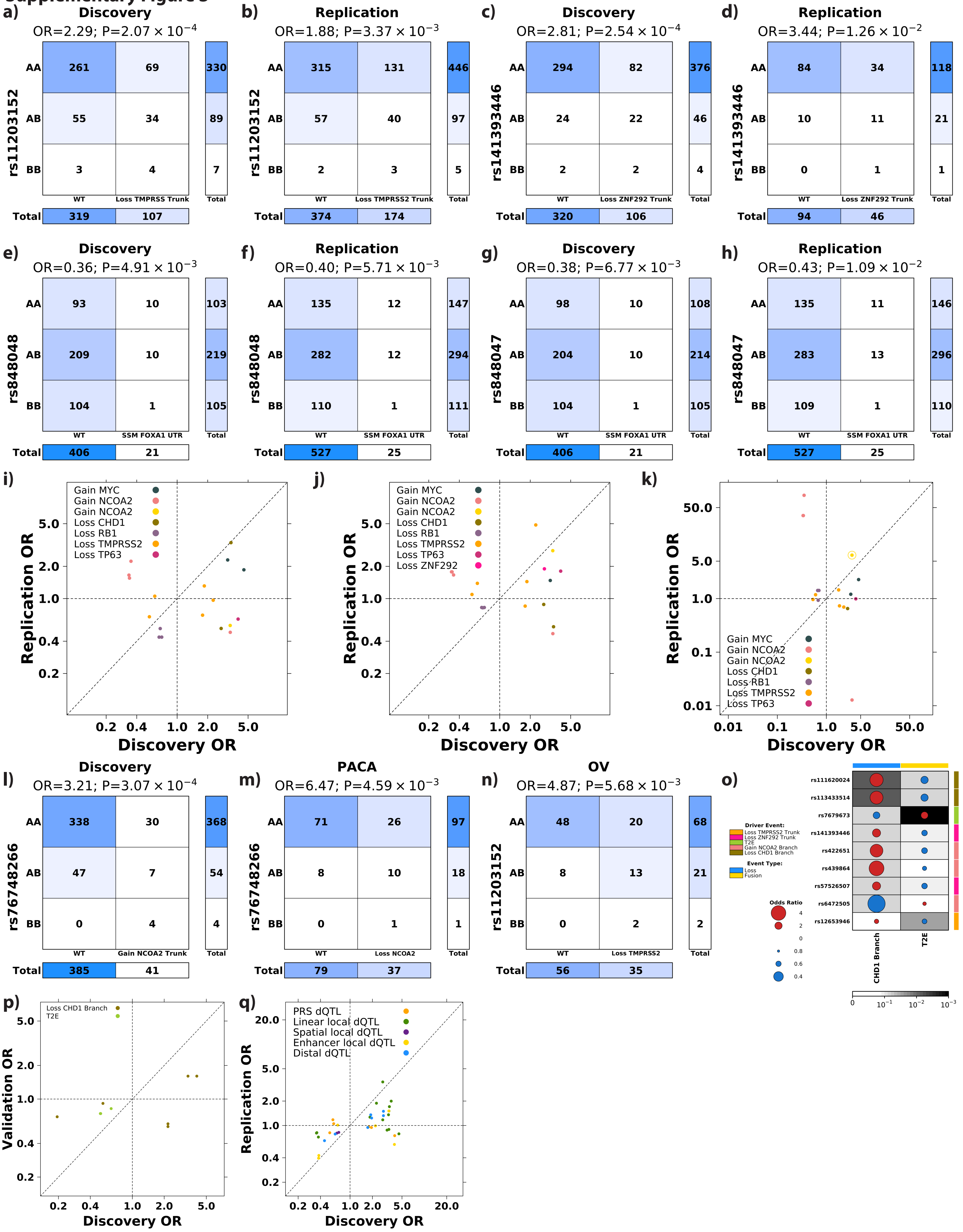

#### Supplementary Figure 3 – Replication of dQTLs

**a-b)** Contingency tables of rs11203152 associated with clonal loss of *TMPRSS2* in discovery cohort **(a)** and replication **(b)** cohort. **c-d)** Contingency tables of rs141393446 associated with clonal loss of *ZNF292* in the discovery **(c)** and replication **(d)** cohorts. **e-f)** Contingency tables of rs848048 associated with SNVs in *FOXA1* 3' UTR in discovery cohort **(e)** and replication **(f)** cohort. **g-h)** Contingency tables of rs848047 associated with SNVs in 3' UTR of *FOXA1* in discovery **(g)** and replication **(h)** cohorts. **i-k)** Comparison of ORs in discovery, x-axis, *vs.* breast **(i)**, ovarian **(j)** or pancreatic **(k)** cancer, y-axis. Only testing dQTLs involving somatic drivers with recurrence rate >5% in each cancer type. Horizontal and vertical dotted lines represent OR = 1 and diagonal line represents y=x. Halo indicates statistical significance (BH FDR < 0.1). **l-m)** Contingency tables of rs76748266 associated with clonal loss of *NCOA2* in the discovery **(l)** and pancreatic **(m)** cohorts. **n)** Contingency table of rs11203152 associated with loss of *TMPRSS2* in ovarian cancer. **o)** Candidate distal dQTLs considering 16 dQTLs with same direction of effect in discovery and replication. **p)** Comparison of ORs in discovery, x-axis, *vs.* replication y-axis, cohorts of candidate distal dQTLs. Dot size and color indicates magnitude and direction of ORs between SNP, y-axis, and driver, x-axis. Covariate along the top indicates the type of somatic event. Covariate along the right indicates the associated somatic driver identified in discovery. **q)** Comparison of ORs in discovery, x-axis, and replication, y-axis, cohorts for 35 dQTLs. Dot color represents strategy used to discovery dQTL. Horizontal and vertical dotted lines represent OR = 1 and diagonal line represents y=x.

### Supplementary Figure 4

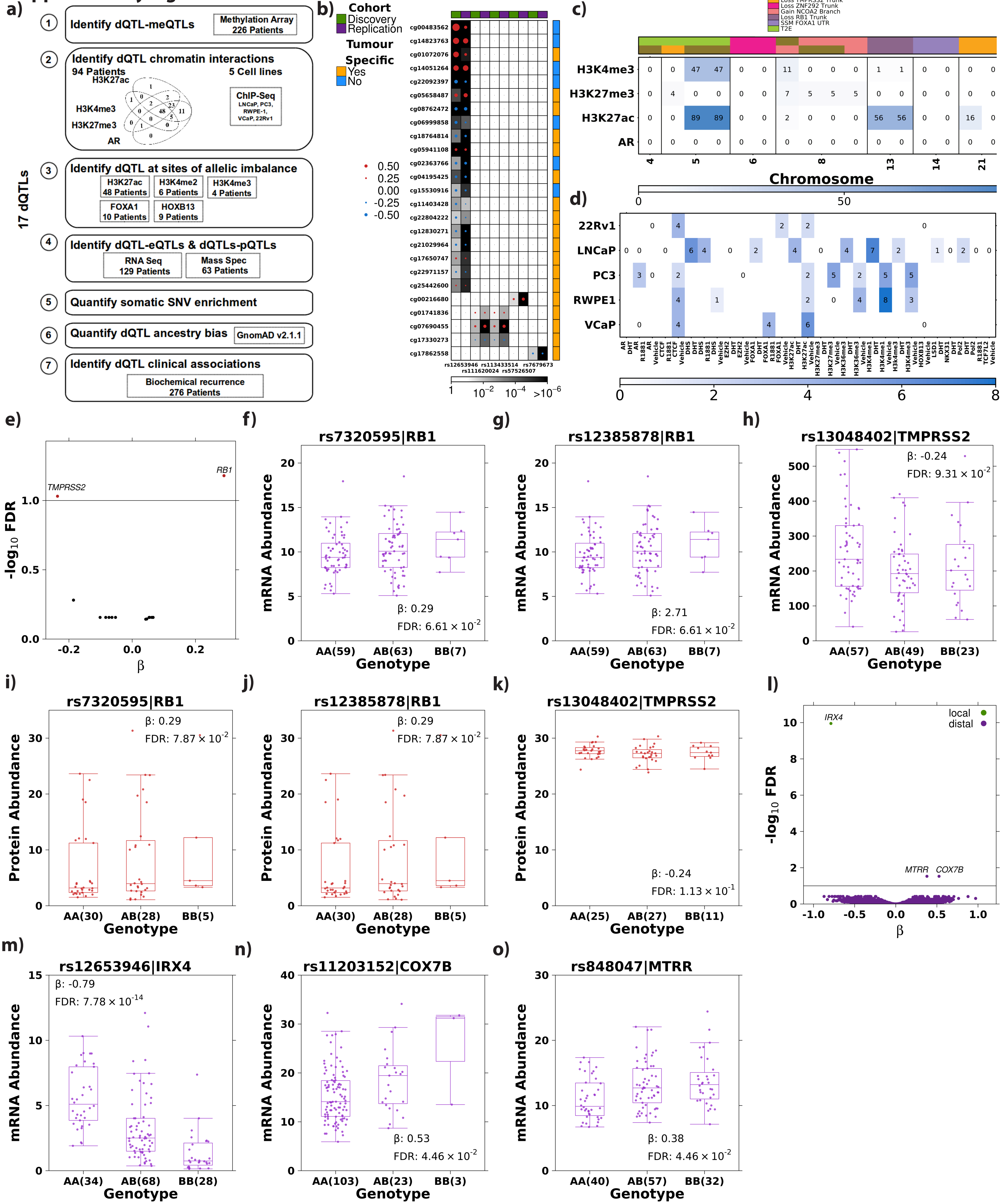

**Supplementary Figure 4 – Molecular characterization of dQTLs**
**a)** Schematic of characterization of dQTLs. **b)** Summary of dQTL-meQTLs. Circle size and color represents effect magnitude and direction of meQTL in discovery and replication cohorts. Background shading indicates false discovery rate. Covariate along the top indicates discovery or replication cohort. Covariate along the right indicates if the meQTL was identified as tumor specific. Highly correlated probes (Spearman's $\rho > 0.80$) are summarized by a single probe and only the top 20 most correlated probes for rs12653946 are plotted. **c)** Overlap of dQTLs, x-axis, with histone modifications and AR binding in primary patient samples, y-axis. Shading indicates the number of patients with overlap. Covariate along the top indicates the somatic driver(s) each SNP is associated with. **d)** Overlap of dQTLs with histone modification and transcription factor binding sites, x-axis, in five prostate cell lines, y-axis. Shading indicates the number of dQTLs that overlap each target in each cell line. X-axis labels give the target and treatment. **e)** Volcano plot of candidate eQTLs – dQTL variant associated with mRNA abundance of associated driver gene. Y-axis shows $\log_{10}$ false discovery rate while x-axis shows effect size of association. Horizontal line indicates BH FDR = 0.1 and red dots indicates a significant association (BH FDR < 0.1). **f-k)** dQTLs are associated with mRNA **(f-h)** and protein **(i-k)** abundance changes of associated driver gene. Boxplot shows mRNA (purple) or protein (red) abundance for gene in title stratified by genotype, x-axis, of the SNP indicated in the title. Statistics are from inverse rank-normalized linear regression model correcting for the first five principal components and age. The number of samples with each genotype is indicated in parenthesis next to the genotype along the x-axis. Boxplot represents median, 0.25 and 0.75 quantiles with whiskers at 1.5x interquartile range. **l)** Volcano plot of local (green) and distal (purple) eQTLs. **m)** One local eQTL identified between rs12653946 and *IRX4*. **n-o)** Two distal eQTLs identified: rs11203152-*COX7B* and rs848047-*MTRR*.

#### Supplementary Figure 5

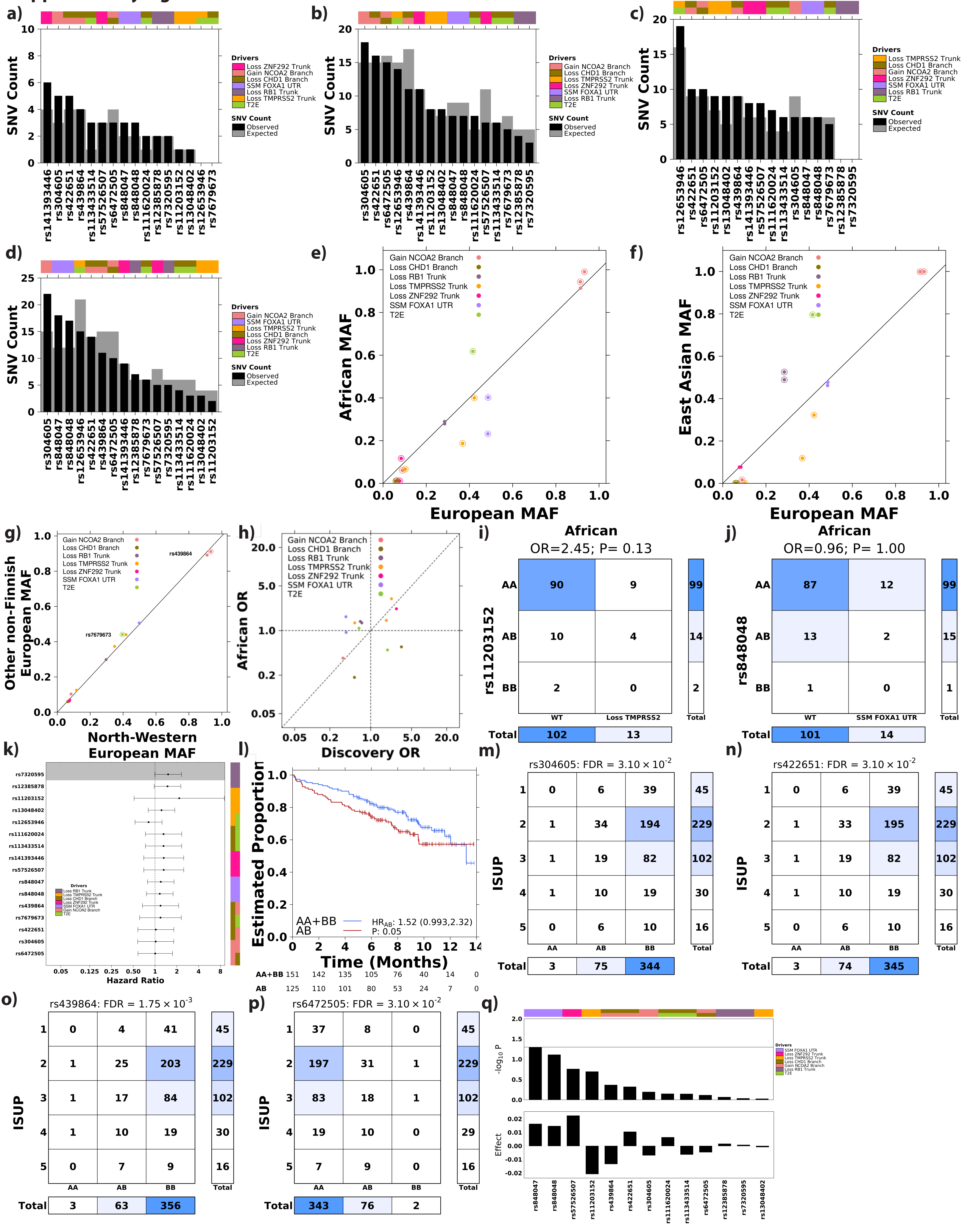

#### Supplementary Figure 5 – Clinical characterization of dQTLs

**a-d)** Number of somatic SNVs within ±10 kbp, y-axis, of each dQTL, x-axis, in prostate **(a)**, breast **(b)**, ovarian **(c)** and pancreatic **(d)** cancer. Background shading indicates number of proximal somatic SNVs expected by chance. Covariate along the top indicates the somatic driver event each SNP is associated with. **e-g)** Comparison of allelic frequencies for 16 dQTLs in European, x-axis *vs.* African, y-axis, populations **(e)**, European *vs.* East Asian populations **(f)** or within European populations **(g)**. Halo indicates SNP has significantly different allele frequencies in two populations. **h)** Comparison of ORs in discovery, x-axis, *vs.* African-descent cohort, y-axis. Horizontal and vertical dotted lines represent OR = 1 and diagonal line represents y=x. **i)** Contingency table of rs11203152 associated with loss of *TMPRSS2* in 115 African men. **j)** Contingency table of rs848048 associated with SNVs in *FOXA1* UTR in 115 African men. **k)** Forest plot showing Hazard Ratios, x-axis, from survival analysis of dQTLs, y-axis, with biochemical recurrence. Error bars represent 95% confidence intervals. Vertical dotted line represents HR = 1. Background shading indicates $P < 0.05$ and covariate on the right indicates the somatic driver event the SNP is associated with. **l)** Kaplan-Meier plot of rs7320595 associated with biochemical recurrence. **m-p)** Contingency tables of association between rs304605 **(m)**, rs422651 **(n)**, rs439864 **(o)** and rs6472505 **(p)** and ISUP grade group. BH FDR from ordinal linear regression. **q)** Barplot shows effect size and p-value from prostate cancer GWAS[9] for 13 non-risk dQTLs, x-axis, with summary statistics from GWAS. Horizontal line indicates P= 0.05. Covariate along the top indicates associated somatic driver.

**Supplementary Figure 6**

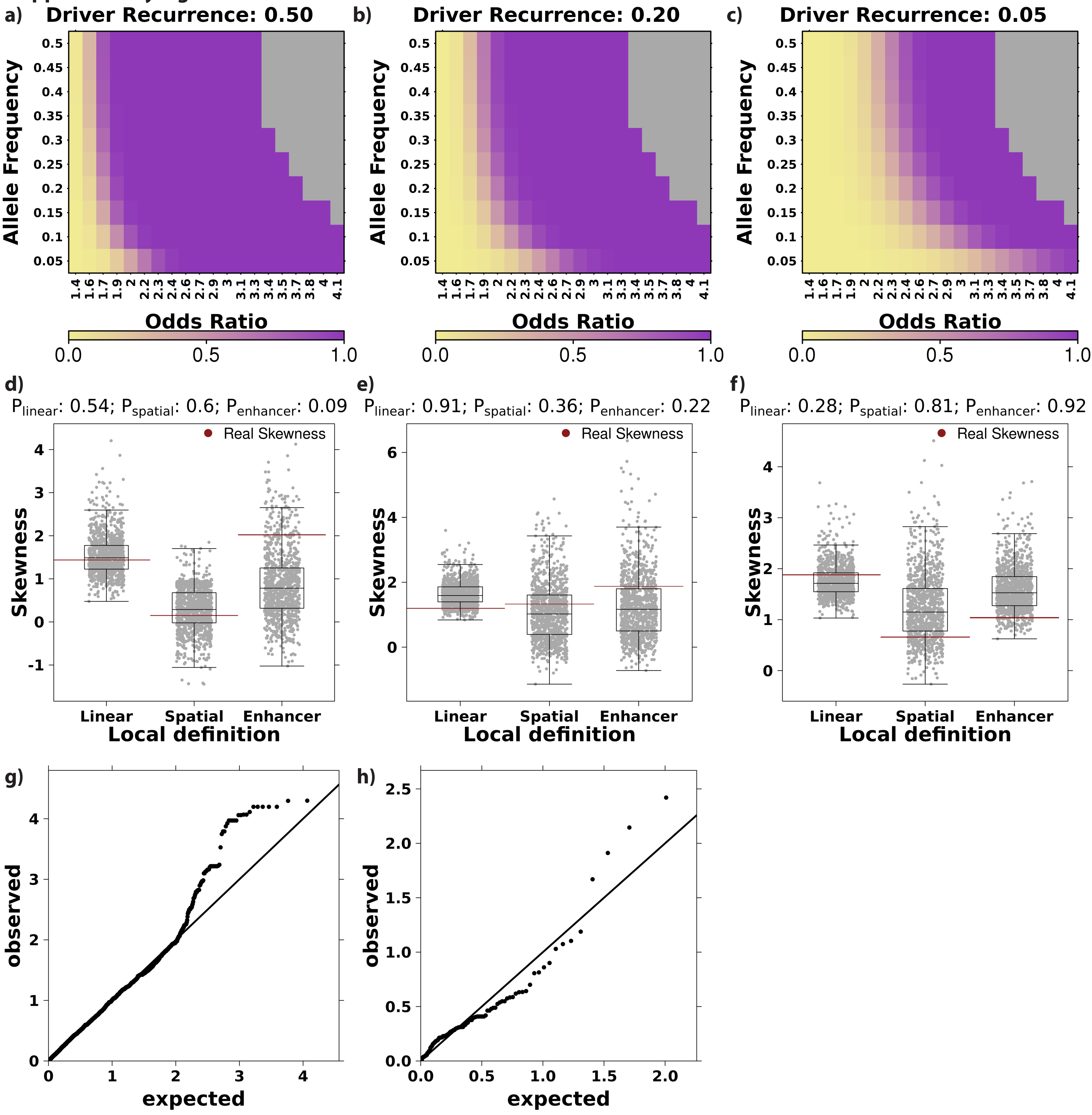

**Supplementary Figure 6 – Enrichment of sub-significance threshold dQTLs**

**a-c)** Heatmaps displaying estimated power considering increasing ORs (x-axis) and allele frequencies (y-axis) for somatic driver recurrence = 0.50 **(a)**, 0.20 **(b)** and 0.05 **(c)**. Shading indicates estimated power with yellow = 0 and purple = 1. Grey indicates power could not be calculated. **d)** Null skew distribution for clonal loss of *RB1* dQTL discovery from 1,000 iterations. Horizontal red lines represent real skew values for each dQTL approach. P-values along the top represent the number of null iterations with skew > real skew divided by the number of null iterations. **e)** Null skew distribution of clonal loss of *NKX3-1*. **f)** Null skew distribution of clonal loss of *TMPRSS2*. **g-h)** Q-Q plots of T2E linear local dQTLs **(g)** and clonal loss of *ZNF292* spatial local dQTLs **(h)**.

### Supplementary Tables

**Supplementary Table 1 – Clinical cohort characteristics and definitions of somatic drivers**

**Supplementary Table 2 – PRS somatic associations**
Summary statistics from PRS and HOXB13 associated with somatic drivers. β and p-value from logistic regression correcting for five genetic principal components, age and somatic mutation burden. BH FDR = Benjamini-Hochberg false discovery rate.

**Supplementary Table 3 – dQTL summary**
Number of dQTLs identified for each somatic driver in each analysis strategy.

**Supplementary Table 4 – Local dQTLs**
Summary statistics from local dQTL associations. Results are not pruned for LD. Statistics from logistic regression correcting for five genetic principal components, age and somatic mutation burden. OR = odds ratio; SE = standard error; L95 = lower 95% confidence interval; U95 = upper 95% confidence interval

**Supplementary Table 5 – dQTL summary statistics across cohorts**
Summary statistics of concordant 23 dQTLs across cohorts

**Supplementary Table 6 – Distal dQTLs**
Summary statistics from distal dQTL associations.

**Supplementary Table 7 – Characterization of dQTLs**
Summary of characterization of concordant 23 dQTLs.

**Supplementary Table 8 – GTEx eQTLs**
dQTL SNPs that were identified as eQTLs in prostate tissue in GTEx.
